# *mRNAbow*: A versatile gene expression system for multiplexed fluorescent imaging using optimized *in vitro* transcribed mRNA

**DOI:** 10.64898/2026.01.02.697412

**Authors:** Heejun Choi, Coral Halanych, William Kasberg, Michael D. Testa, Sarah Rubin-Elgressy, Phuong Nguyen, Deepika Walpita, Arthur Tsang, Dan Cortes, Erin Yue Song, Haodi Wu, Irving L Weissman, Isabel Espinosa-Medina, Chie Satou, Jia L. Song, David Q. Matus, Jennifer Lippincott-Schwartz

## Abstract

Messenger RNA (mRNA) transfection enables rapid, transient protein expression without nuclear entry, providing a powerful alternative to DNA or viral delivery in post-mitotic and otherwise difficult-to-transfect cells. Although *in vitro* transcribed (IVT) mRNAs have revolutionized therapeutic applications, their adoption in experimental biology remains limited by challenges in synthesis, variability across cell types, and concerns about cytotoxicity. Here, we define design principles that maximize IVT mRNA performance across diverse cellular and organismal systems. Through systematic comparison of capping strategies and base modifications, including N1-methyl-pseudouridine, 5-methylcytidine, and 5-methoxyuridine, we identify modifications that enhance translation while minimizing activation of cellular stress responses. Optimized transcripts drive robust protein expression within four hours, persist for up to one week, and support multiplexed expression of structurally and functionally distinct proteins in mammalian cells, including cancer cell lines, iPSC-derived systems, primary cells, and organoids, as well as *in vivo* in zebrafish embryos and in less genetically tractable models such as *Danionella cerebrum* and sea urchin embryos. To further expand accessibility for community use, we developed *mRNAbow*, a platform for generating low-toxicity mRNAs encoding organelle-targeted fluorescent proteins and biosensors for multiplex imaging, with corresponding plasmids made publicly available. Together, these advances establish a generalizable framework for IVT mRNA design and expand experimental access to synthetic mRNA technologies for dissecting cellular architecture and dynamics.

## Introduction

Messenger RNA (mRNA) transfection offers a powerful approach for transient protein expression in mammalian cells. Unlike plasmid DNA, synthetic mRNAs do not require nuclear entry and are translated immediately in the cytoplasm, enabling efficient expression in post-mitotic and other hard-to-transfect cell types, including neurons, hepatocytes, and macrophages^1,2^. These advantages have underpinned the clinical success of *in vitro* transcribed (IVT) mRNAs, most notably for mRNA vaccines^3^ and cancer therapeutics^4^. In contrast, viral delivery methods often show poor infectivity or frequently induce strong antiviral responses in immune cells and other primary cell types^5^, limiting their usefulness for achieving controlled and uniform transgene expression. Despite their promise, IVT mRNAs are still not widely used as routine experimental tools in biology, partly due to practical challenges to synthesis, earlier concerns about cytotoxicity^6^, and the lack of accessible reagent libraries for efficient construct design.

Studies in mRNA therapeutics established that carefully engineered untranslated regions (UTRs)^7,8^, improved co-transcriptional capping chemistries^9,10^, and the incorporation of modified nucleotides such as N1-methyl-pseudouridine (m1Ψ) or 5-methylcytidine (m5C) can substantially enhance protein yield while mitigating stress responses^6^. Despite these advances, systematic evaluation of these design parameters across diverse biological contexts remains limited, particularly in the context of cell biology applications. Modified nucleotides are often assumed to uniformly improve translation efficiency and suppress innate immune activation, yet their effects vary between cell types and can involve trade-offs in translation efficiency^11,12^, RNA stability^13,14^, or protein synthesis fidelity^15,16^. Developing broadly applicable methods for IVT mRNA production and delivery is therefore essential to make these reagents accessible for experimental biology. This need is especially pressing in systems like primary cultures and tissues, where biosafety constraints, low transfection efficiency, and short experimental windows limit throughput.

Here, we demonstrate IVT mRNAs can be used to express a wide range of proteins and biosensors in single cells, tissues or whole organisms, overcoming challenges of low protein expression levels, cytotoxicity or mRNA stability. We first evaluated the impact of capping strategies and base modifications on mRNA transfection outcomes across multiple mammalian cell types and *in vivo* during zebrafish embryogenesis. Using a standardized reporter construct, we compared unmodified transcripts with those incorporating m1Ψ, m5C, both, or 5-methoxyuridine (5moU), with and without of co-transcriptional capping. Our analyses showed that capping is required for efficient protein expression and revealed cell-type-specific preferences for distinct nucleotide modifications. Live-cell imaging detected protein production within four hours of transfection, with expression scaling in a dose-dependent manner and persisting for 48 hours to one week. The optimized transcripts supported robust expression of structural proteins, biosensors, and organelle markers. Furthermore, simultaneous delivery of multiple mRNAs enabled multiplexed labeling of different organelles within single cells. To promote broader access, we generated a palette of mRNAs encoding structural, organellar, and sensor proteins (*mRNAbow*), made publicly available as plasmids in the *4DCP mRNAbow Collection*. Together, these findings establish a generalizable framework for IVT mRNA design and application, providing a practical toolkit for cell biology.

## Results

### Chemical modifications and capping in IVT mRNA expression

To establish a robust template for *in vitro* transcribed (IVT) mRNAs, we incorporated well-characterized regulatory elements known to enhance translation efficiency (**Fig. 1A**). Specifically, optimized 5’ and 3’ untranslated regions (UTRs) were used as these have been shown to maximize protein output upon transfection^8^. A ∼50-adenosine stretch was included to generate a poly(A) tail, and the entire expression cassette was cloned into a plasmid backbone containing a T7 promoter. This configuration enabled co-transcriptional addition of a 5’ cap structure using CleanCap M6 technology, which generates a Cap1 structure and protects exogenous IVT mRNA from decapping^17,18^.

**Figure 1.**
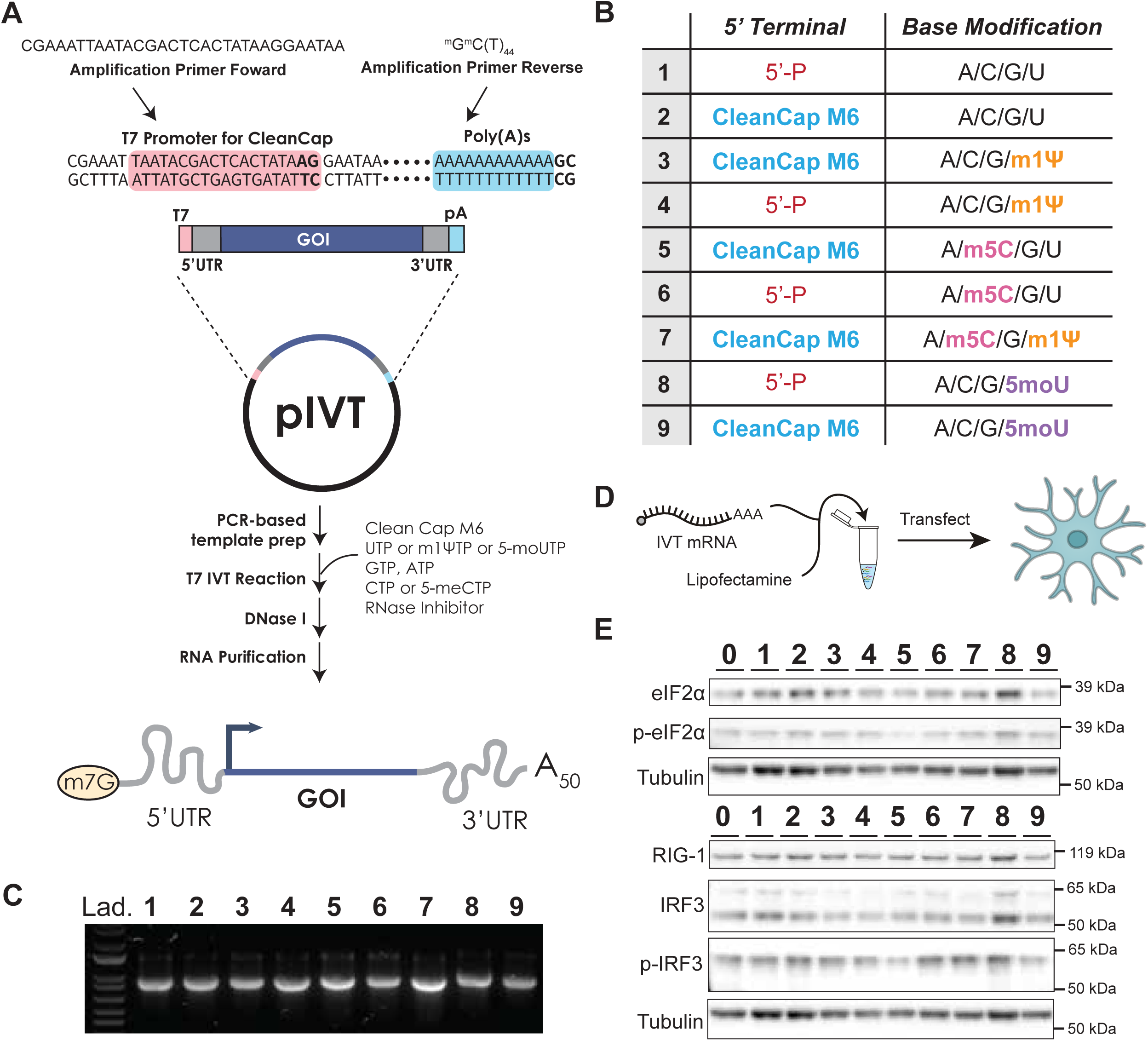
Design, synthesis, and cellular delivery of modified IVT mRNAs without activation of innate stress responses. **A**, Schematic of IVT mRNA construct design. The template contains optimized 5’ and 3’ UTRs flanking a coding sequence, followed by a 50-adenosine poly(A) tail. Linearized templates are generated by PCR using a forward primer annealing to the T7 promoter and a reverse primer containing two 2’-O-methylated ribonucleotides at the 5’ end to minimize dsRNA byproducts. *In vitro* transcription is performed with mutant T7 polymerase in the presence of CleanCap M6 for co-transcriptional capping along with different NTPs. **B**, Summary of mRNA constructs tested. Nine variants were generated incorporating different combinations of 5’ cap status (uncapped or CleanCap M6) and base modifications: unmodified (constructs **1, 2**), N1-methylpseudouridine (m1Ψ; constructs **3, 4**), 5-methylcytidine (m5C; constructs **5, 6**), m1Ψ + m5C (construct **7**), or 5-methoxyuridine (5moU; constructs **8, 9**). **C,** Agarose gel electrophoresis of purified IVT mRNAs encoding nuclear mScarlet3 showing high yield and purity for all nine constructs. **D,** Schematic illustrating IVT mRNA transfection. IVT mRNAs were complexed with lipofectamine to form lipid nanoparticles, which were then delivered into cells to enable transient protein expression. **E,** Immunoblot images of stress response markers in U-2 OS cells 24 hours post-transfection. Lanes correspond to non-transfected control (**0**) and constructs **1-9**. No induction of phosphorylated eIF2α (p-eIF2α), RIG-I, or phosphorylated IRF3 (p-IRF3) was detected across conditions, indicating absence of dsRNA-triggered stress responses. Total IRF3, eIF2α, and tubulin serve as loading controls.

To prepare linearized DNA templates, we designed a forward primer annealing to the T7 promoter and a reverse primer containing two 2’-O-methylated ribonucleotides. These terminal modifications have been reported to reduce double-stranded RNA (dsRNA) byproducts during transcription^19^. The cleaned PCR product was then used for *in vitro* transcription with T7 polymerase in the presence of the appropriate rNTPs. A mutant T7 polymerase was employed to further minimize dsRNA formation. Detailed procedures for mRNA synthesis are provided in the **Supplementary Information**.

We next generated a panel of mRNAs incorporating defined chemical base modifications and distinct 5’ capping variants (**Fig. 1B**) to systematically assess their effects on transfection efficiency and resulting protein expression. For streamlined quantification, we used a nuclear-targeted mScarlet3 construct. Capped or uncapped mRNAs were synthesized incorporating either unmodified bases (constructs **1**, **2**), N1-methylpseudouridine (m1Ψ; constructs **3**, **4**), 5-methylcytidine (m5C; constructs **5**, **6**), a combination of m1Ψ and m5C (construct **7**), or 5-methoxyuridine (5moU; constructs **8**, **9**), each with or without CleanCap M6 (**Fig. 1B**). All variants were transcribed at high yield and purity, as confirmed by agarose gel electrophoresis (**Fig. 1C**).

Purified mRNAs were complexed with lipofectamine and transfected into U-2 OS cells (**Fig. 1D**). Transfection of nuclear mScarlet3 mRNA into U-2 OS cells did not trigger detectable cellular stress responses typically associated with double-stranded RNA accumulation (**Fig. 1E)**. In particular, we observed no activation of integrated stress response such as eIF2α phosphorylation, nor dsRNA-specific stress response markers such as RIG-I expression or IRF3 phosphorylation, relative to non-transfected controls (denoted as **0**) (**Fig. 1E** and **Supplementary Figure 1**).

Transfection of 100 ng nuclear mScarlet3 mRNA into U-2 OS cells (approximately 30,000 cells per well) yielded robust nuclear fluorescence within 24 hours (**Fig. 2A**). Fluorescence microscopy revealed strong, uniform nuclear signal across most transfected cells. Using nuclear DNA staining to define each nucleus, we quantified both the fraction of mScarlet3-positive nuclei (transfection efficiency, **Fig. 2B**) and the mean nuclear fluorescence intensity as a measure of protein expression level (**Fig. 2C**).

**Figure 2.**
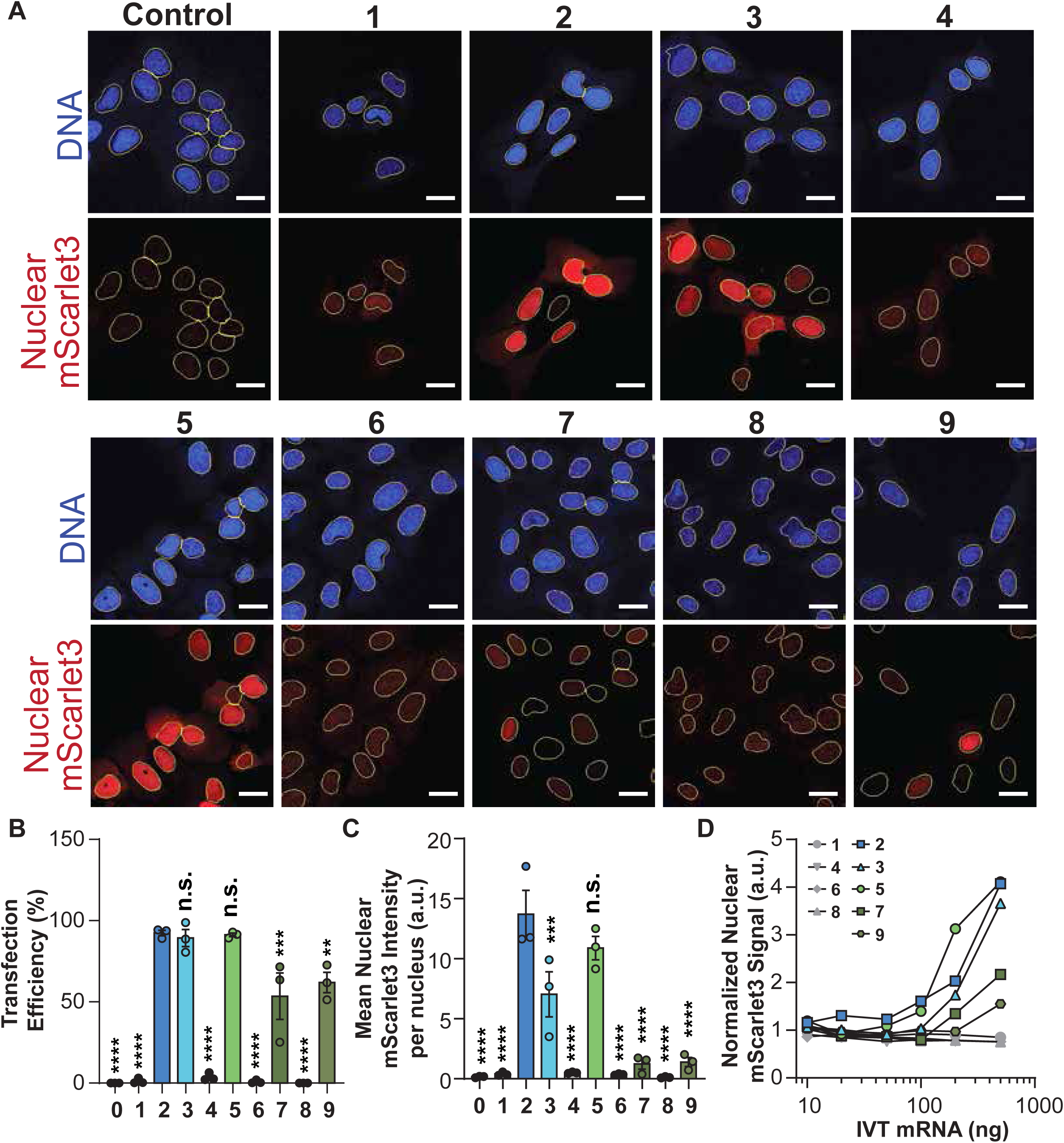
Effects of nucleotide modification and capping on IVT mRNA expression in U-2 OS cells. A, Representative fluorescence microscopy images of U-2 OS cells transfected with 100 ng of nuclear mScarlet3 mRNA variants (constructs 1-9). Blue: Hoechst nuclear stain; Red: mScarlet3. Scale bar, 10 µm. **B**, Quantification of mean nuclear mScarlet3 fluorescence intensity per cell for each construct. Data are normalized to capped unmodified mRNA (construct **2**, set to 100%). Capped unmodified mRNA yielded the highest expression, while m5C incorporation (construct **5**) reduced expression by ∼20%, m1Ψ (construct **3**) by ∼49%, and m1Ψ + m5C (construct **7**) and 5moU (construct **9**) showed >85% reduction. Uncapped constructs (**1, 4, 6, 8**) showed negligible signal. Data represent mean ± SEM from n = 3 independent experiments. Statistical tests were performed against construct **2**. **C**, Transfection efficiency calculated as percentage of mScarlet3-positive nuclei. Capped mRNAs with unmodified bases (construct **2**), m1Ψ (construct **3**), or m5C (construct **5**) achieved >89% efficiency, while combined modifications (construct **7**) and 5moU (construct **9**) showed reduced efficiency (∼54% and ∼62%, respectively). Uncapped constructs yielded minimal transfection. Data represent mean ± SEM from n = 3 independent experiments. Dunnett’s test was performed against construct **2**. **D**, Dose-response relationship showing mean nuclear fluorescence intensity as a function of transfected mRNA amount.

Among all tested conditions, capped unmodified mRNAs produced the highest fluorescence intensity and transfection efficiency (**Fig. 2B** and **2C**, construct **2**, 92.4%). As expected, uncapped mRNAs failed to support translation, confirming that cap addition is essential, although the absence of a cap (constructs **1**, **4**, **6**, **8**) or chemical modification (**2**) did not trigger a stress response (**Fig. 1E**). m5C incorporation moderately reduced expression (**Fig. 2C**, construct **5**, 20% lower than capped unmodified mRNAs) while maintaining high transfection efficiency (**Fig. 2B**, construct **5**, 91.3%), whereas m1Ψ incorporation further lowered signal intensity (**Fig. 2C**, construct **3**, 49% lower than capped unmodified mRNAs) while still achieving high transfection efficiency (**Fig. 2B**, construct **3**, 89.4%). In contrast, 5moU incorporation and the combination of m1Ψ and m5C resulted in the weakest fluorescence (**Fig. 2C**, constructs **9** and **7**, 89.8% and 89% reduction compared to capped unmodified mRNAs, respectively), consistent with reduced transfection efficiency (**Fig. 1B**, constructs **9** and **7**, 61.9% and 54%, respectively).

Together, these results demonstrate that while base modifications can markedly influence the translational output of synthetic mRNAs, capped unmodified transcripts (construct **2**) remain the most effective for robust protein expression in U-2 OS cells, with m5C (construct **5**) and m1 Ψ (construct **3**) showing moderately reduced but still efficient expression. Expression levels scaled proportionally with the amount of transfected mRNA (**Fig. 2D**), demonstrating clear dose dependency. This property makes the system highly adaptable, allowing users to modulate expression level by varying mRNA input and modifications.

### Temporal dynamics of mRNA delivery and protein production

After characterizing how chemical modifications influence mRNA expression in U-2 OS cells, we next examined the temporal dynamics of mRNA delivery and subsequent protein synthesis. We used quantitative RT-PCR to measure intracellular delivery and stability by quantifying nuclear-targeted mScarlet3 transcripts. For both uncapped (**1**) and capped transcripts (**2, 3, 5, 7 and 9**), cytoplasmic mRNA level increased progressively during the first 8 hours after transfection, then plateaued and remained stable for over 24 hours (**Fig. 3A**). These results demonstrate that uptake and cytoplasmic stability of exogenous mRNAs occur largely independently of specific base modifications or 5’ cap structures. Because mRNA stability was similar across all variants, whereas protein expression differed substantially, variation in translational efficiency likely accounts for the observed differences in protein output seen in **Fig. 2C**.

**Figure 3.**
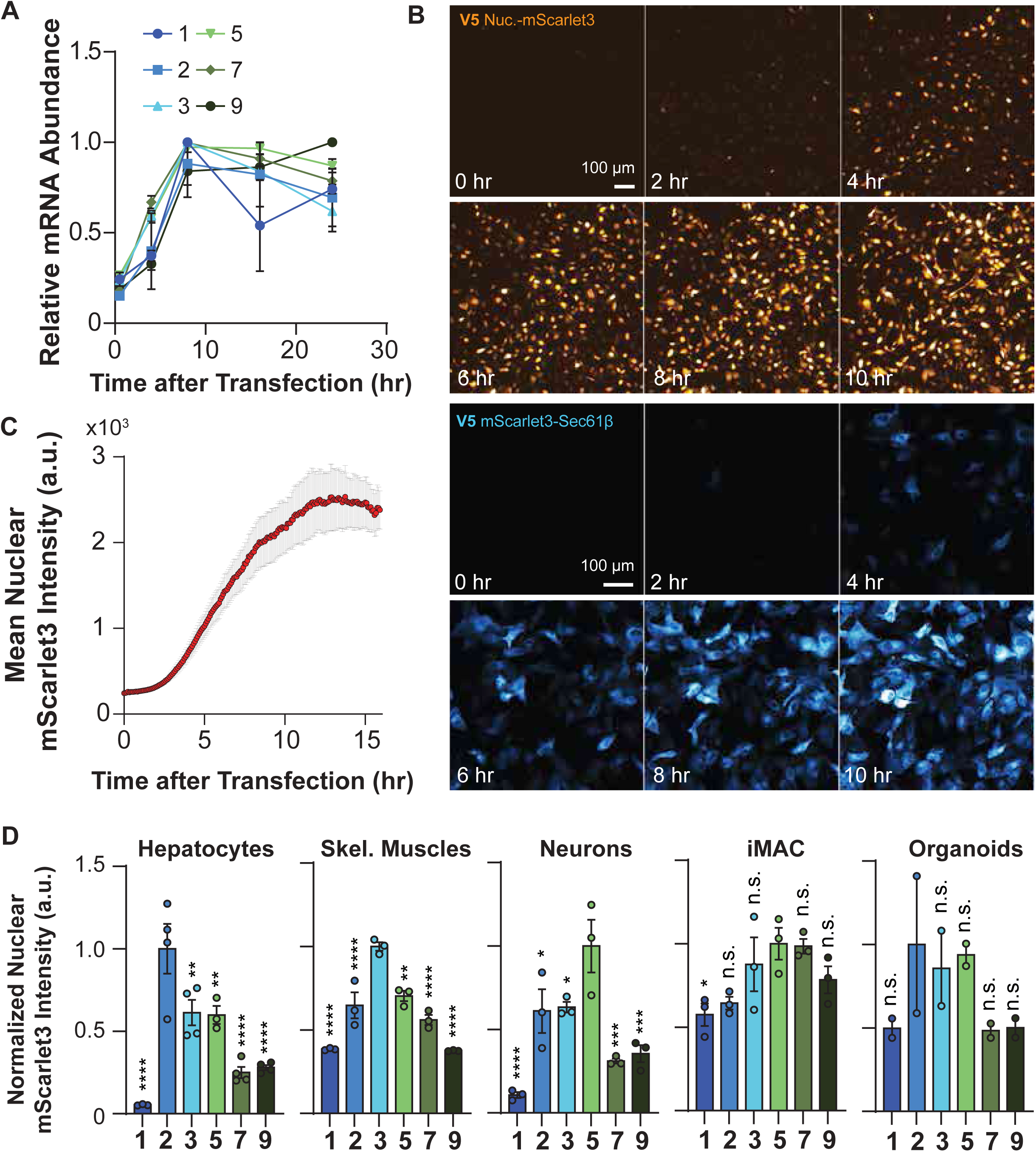
mRNA stability, protein expression kinetics, and cross-cell type performance of IVT mRNAs. A, Relative mRNA abundance measured by RT-qPCR time course of intracellular nuclear mScarlet3 mRNA levels in U-2 OS cells following transfection with constructs **1, 2, 3, 5, 7,** and **9**. Data are normalized for each construct. Mean ± SEM from n = 3 independent experiments. **B**, Live-cell imaging time course of protein production in U-2 OS cells transfected with capped m5C-modified mRNA encoding nuclear-targeted mScarlet3 (orange, top) or mScarlet3-Sec61β (ER marker, Cyan, Bottom) in U-2 OS cells. Representative images shown at indicated timepoints. Scale bars, 100 µm. **C**, Nuclear mScarlet3 intensity per nucleus following mRNA transfection. The line represents the 95% confidence interval.

To monitor protein production dynamics directly, we performed live-cell imaging using capped m5C-modified mRNAs encoding nuclear-targeted mScarlet3 and mScarlet3-Sec61β over an 18-hour period. Detectable fluorescence appeared within 4 hours of transfection (**Fig. 3B, 3C, Video S1 and S2**), continued to increase, and reached a plateau around 16 hours. These kinetics highlight the rapid onset of protein production enabled by this delivery strategy. The plateau in mRNA abundance measured by qPCR at 8 hours likely reflects the completion of lipid-mediated delivery, whereas sustained protein expression beyond this timepoint indicates that translation continues efficiently after mRNA internalization has stabilized.

### Primary cells show cell-type-specific modification preferences

We next examined whether the trends observed in U-2 OS cells extended to other biological systems. To do this, we evaluated mRNA transfection efficiency and expression across a range of multiple primary and differentiated cell types, including adult mouse hepatocytes, rat hippocampal neurons, human skeletal muscle cells, iPSC-derived macrophages, and iPSC-derived brain organoids. These post-mitotic and primary cells are typically resistant to DNA-based transfection methods, making mRNA-based approaches particularly valuable. Under the similar transfection conditions comparable to those used for U-2 OS cells, robust protein expression required a 5’ cap in all cell types tested (**Fig. 4A**; uncapped construct **1** vs. capped construct **2, Supplementary Fig. 2**), confirming the universal importance of cap structure in mediating efficient translation.

**Figure 4.**
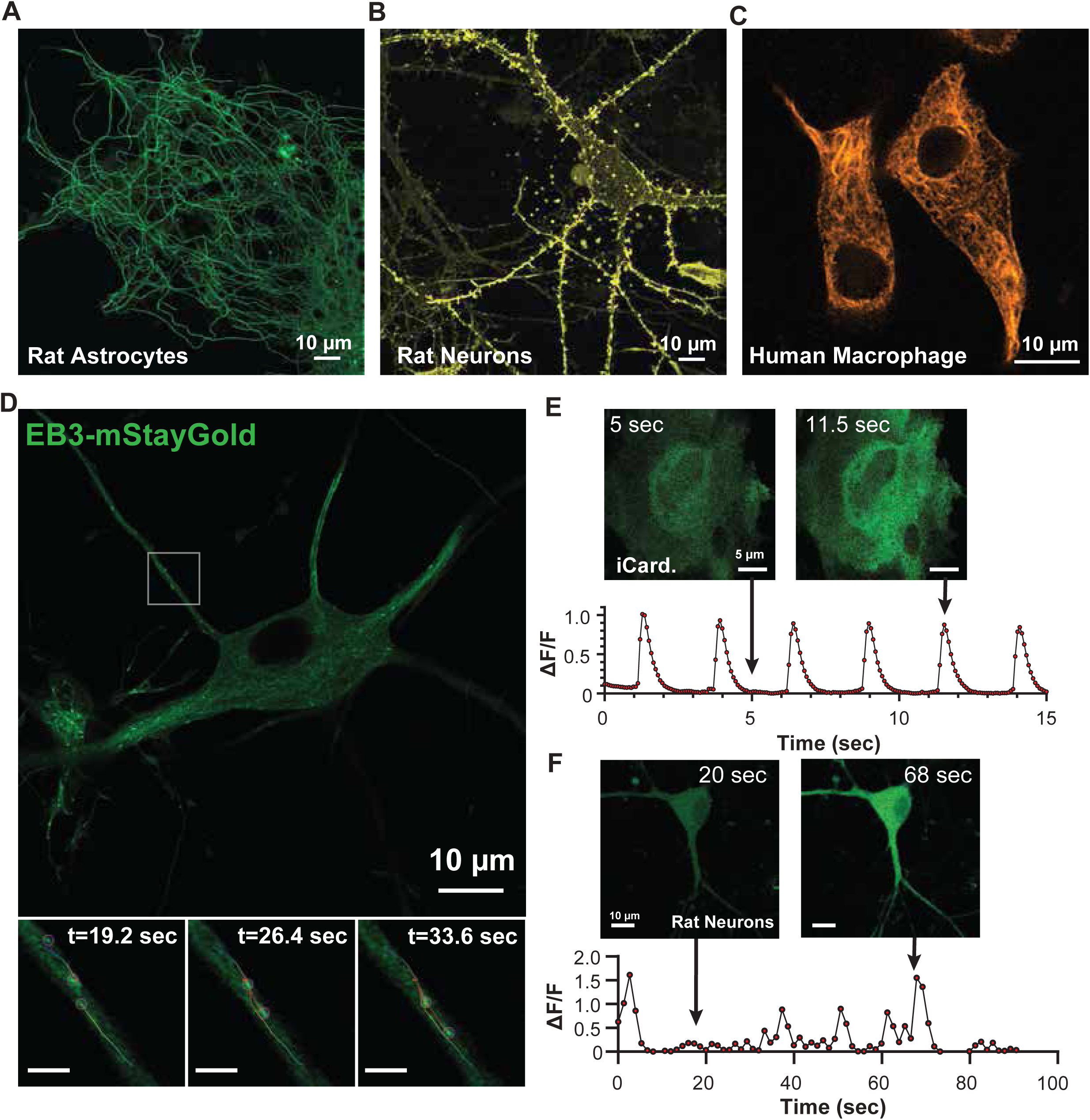
Broad applicability of IVT mRNAs for structural and functional imaging across primary cell types. A, Comparison of mRNA construct performance across primary and differentiated cell types. Mean fluorescence intensity was quantified 24 hours post-transfection for adult mouse hepatocytes, rat hippocampal neurons, human skeletal muscle cells, iPSC-derived macrophages, and iPSC-derived brain organoids. Mean ± SEM from n = 3 independent experiments for all except iPSC-derived brain organoid (n =2). Dunnett’s test was performed against the highest expressing construct. **B-D**) Representative images of cytoskeletal structures in primary cells transfected with capped m5C-modified mRNAs encoding (**B**) Ensconsin-mStayGold (microtubules) in rat astrocyte, (**C**) LifeAct-mScarlet3 (F-actin) in rat hippocampal neurons, and (**D**) Vimentin-mScarlet3 (intermediate filaments) in human monocyte-derived macrophages. Scale bars, 10 µm. **E,** Dynamic imaging of microtubule plus-ends in rat hippocampal neurons expressing EB3-mStayGold. Top: Representative image showing characteristic comet-like signals in soma and dendrites. Bottom: Trajectory analysis of EB3 comets along a dendrite segment, revealing bidirectional movement and growth reversals. Scale bars, 10 µm (top) and 1 µm (bottom) **F,** Calcium dynamics in iPSC-derived cardiomyocytes expressing GCaMP8s from synthetic mRNA. Top: Images showing low (left) and high (right) calcium transients. Bottom: Fluorescence traces (ΔF/F) from the same cell. Scale bar, 5 µm. **G,** Spontaneous calcium activity in rat hippocampal neurons expressing GCaMP8s. Images showing low (left) and high (right) calcium transients. Bottom: Fluorescence traces (ΔF/F) from the same cell. Scale bar, 10 µm.

Distinct preferences for nucleotide modifications emerged across cell types. Hepatocytes showed highest expression from capped, unmodified mRNAs (**Fig. 4A**, Hepatocytes, construct **2**; **Supplementary Fig. 2A**), mirroring the pattern observed in U-2 OS cells (**Fig. 1G**, construct **2**). In contrast, capped m1Ψ-modified mRNAs produced maximal expression in human skeletal muscle cells (**Fig. 4A**, Skeletal muscle, construct **3**; **Supplementary Fig. 2B**), whereas capped m5C-modified mRNAs yielded stronger fluorescence in rat hippocampal neurons (**Fig. 4A**, Neurons, construct **5**; **Supplementary Fig. 2C**). iPSC-derived macrophages expressed comparably from capped m1Ψ- and m5C-modified mRNAs (**Fig. 4A**, Macrophages, constructs **3, 5,** and **7**; **Supplementary Fig. 2D**), while iPSC-derived brain organoids exhibited similar expression from capped unmodified, m1Ψ-, and m5C-modified mRNAs (**Fig. 4A**, Organoids, constructs **2, 3,** and **5**; **Supplementary Fig. 2E**).

These systematic comparisons of different nucleotide modifications allowed us to identify a modification that could provide reliable performance across systems. Given that recent studies suggest that m1Ψ incorporation can increase translational frameshifting^16^, and given our observation that m5C consistently promoted strong expression across diverse cell types, we selected m5C (construct **5**) as the most reliable modification for subsequent experiments. This choice is particularly advantageous for C-terminally tagged reporters, where frameshifting could result in protein mislocalization or loss of function. Across hepatocytes, neurons, muscle cells, macrophages, and brain organoids, capped mRNAs containing m5C consistently drove strong expression, establishing m5C as a robust, generalizable modification for IVT mRNA expression across diverse cellular contexts

### Synthetic mRNA enables live imaging of cytoskeleton and calcium dynamics

We next tested whether individual mRNA transfections of established reporter constructs could recapitulate their known localization and functional behaviors. As a first step, we synthesized capped mRNAs with m5C modification and an M6 CleanCap that encode structural markers: Ensconsin-mStayGold for microtubules (**Fig. 4B**), LifeAct-mScarlet3 for filamentous actin (**Fig. 4C**), and Vimentin-mScarlet3 for intermediate filaments (**Fig. 4D**). Expression from these transcripts produced bright, well-defined labeling of cytoskeletal components in live primary cultures, consistent with endogenous organization.

To visualize microtubule dynamics, we generated EB3-mStayGold mRNA, encoding the plus-end-binding protein EB3, a classic marker for microtubule growth. Following transfection into primary rat hippocampal neurons, EB3–mStayGold produced the characteristic comet-like signals at microtubule plus ends in both soma and dendrites **(Fig. 4E; Video S3)**. Single-particle tracking of individual comets revealed bidirectional movement and occasional growth reversals along dendrites **(Fig. 4E, bottom panels; Video S4)**, consistent with established microtubule behavior. When expressed in human skeletal muscle cells, EB3–mStayGold displayed strictly unidirectional comet movement, reflecting the uniform microtubule polarity characteristic of myotubes **(Supplementary Fig. 3A; Video S5)**

We then tested whether functional biosensors could be expressed across different primary cell types. We synthesized mRNA expressing GCaMP8s, a highly sensitive genetically encoded calcium indicator optimized for high signal-to-noise ratio^20^, with m5C modification and an M6 cap. The synthesized mRNA was delivered to iPSC-derived cardiomyocytes (iCardiocytes) and rat hippocampal neurons, which exhibit distinct patterns of spontaneous calcium signaling. In iCardiocytes, GCaMP8s reported rhythmic calcium oscillations of uniform amplitude, with each transient reaching a similar ΔF/F (**Fig. 4F**). In addition, smaller calcium sparks often preceded the main transients, consistent with prior reports^21^ (**Fig. 4F** and **Video S6**). In hippocampal neurons, spontaneous firing produced heterogeneous calcium responses, reflected as variable ΔF/F amplitudes, consistent with previously observed viral-transduced GCaMP8s expression in neurons^20^ (**Fig. 4G**). Finally, transfection of 2 µg GCaMP8s mRNA into acute rat brain slices resulted in detectable spontaneous calcium transients, confirming that the biosensor can report neural activity in *ex vivo* tissue **(Video S7)**.

Collectively, these results demonstrate that synthetic mRNAs incorporating m5C modification and an M6 CleanCap enable robust, transient expression of both structural markers and functional biosensors. This approach supports high-quality live imaging of cytoskeletal organization and calcium dynamics across diverse primary cell types and tissue preparations.

### mRNA cocktails enable multiplexed organelle labeling

Unlike viral vectors, which typically deliver a single transgene per particle, lipid-based delivery systems can encapsulate multiple mRNAs simultaneously^22^. We hypothesized that this property could enable efficient co-delivery of distinct reporters to the same cell. To test this, we generated mRNAs encoding nuclear-targeted fluorescent proteins, including mTagBFP2, mStayGold, mScarlet3, and HaloTag, and transfected U-2 OS cells either individually or in combination. The mRNAs were premixed before complexation with the transfection reagent, allowing random co-packaging within lipids and simultaneous delivery of multiple transcripts to single cells (**Fig. 5A**). Co-expression of the different reporters was evident as distinct nuclear-localized fluorescence signals corresponding to mTagBFP2, mStayGold, mScarlet3, and JF646-labeled HaloTag (**Fig. 5B**).

**Figure 5.**
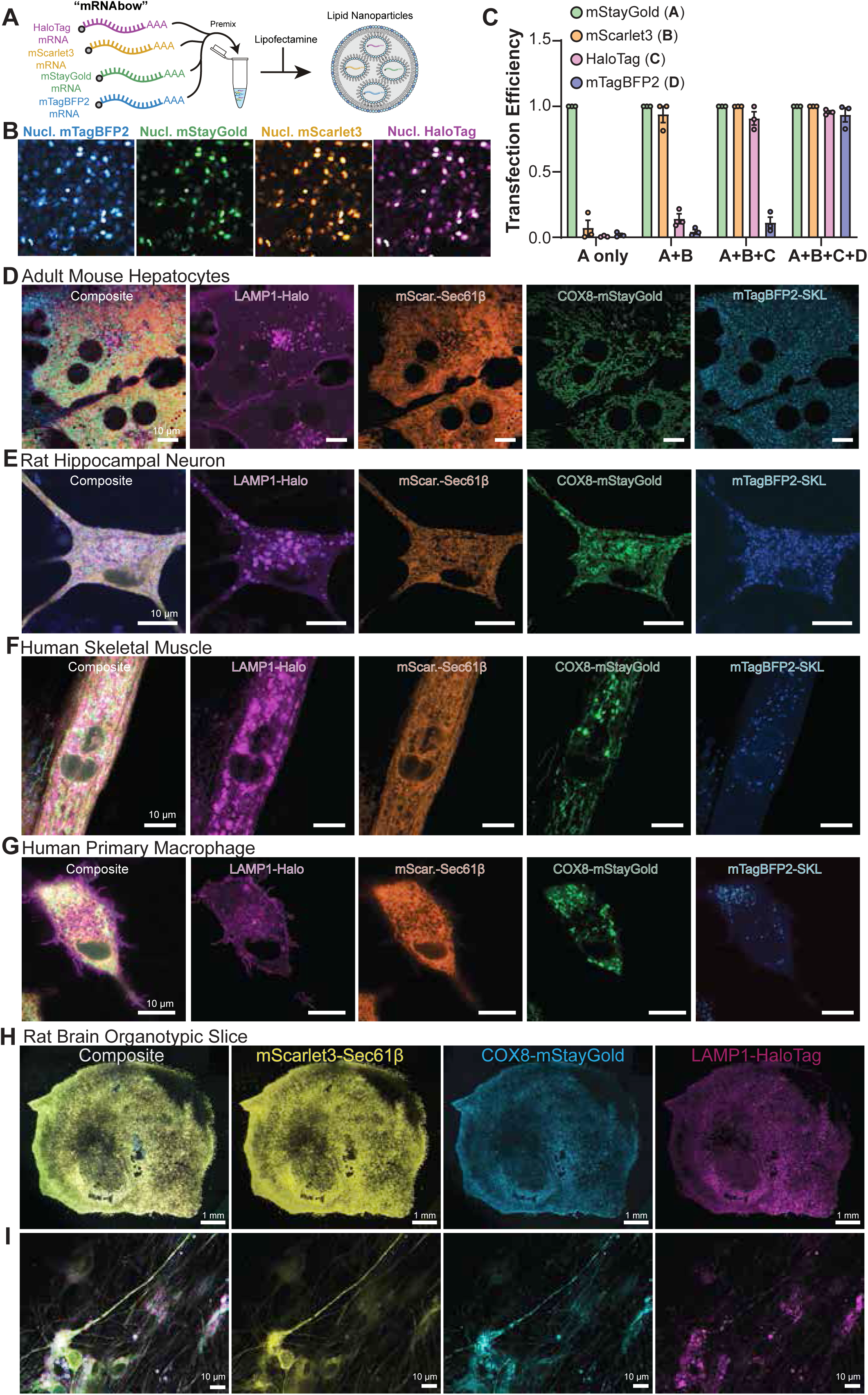
*mRNAbow* enables multiplexed organelle labeling in primary cells and tissue slices. A, Schematic of pre-mixed mRNA cocktails (*mRNAbow*) encoding distinct proteins followed by lipofectamine. **B** Co-expression of four nuclear fluorescent proteins (mTagBFP2, mStayGold, mScarlet3, HaloTag) in U-2 OS cells. Scale bar, 50 µm. **C** Co-transfection efficiency quantified from mStayGold-segmented nuclei (**A**) and co-expression of mScarlet3 (**B**), HaloTag (**C**), and mTagBFP2 (**D**) across one- (**A only**), two- (**A+B**), three- (**A+B+C**), and four-mRNA (**A+B+C+D**) delivery conditions. **D-G** Simultaneous expression of four organelle markers from co-delivered mRNA cocktails (LAMP1-HaloTag for lysosomes, mScarlet3-Sec61β for ER, 4xCOX8-mStayGold for mitochondria, mTagBFP2-SKL for peroxisomes) in diverse primary cell types: **D**, adult mouse hepatocytes, **E**, rat hippocampal neurons, **F**, human skeletal muscle cells, and **G,** human monocyte-derived macrophages. All mRNAs contained m5C modifications and CleanCap M6. Images acquired 24-48 hours post-transfection demonstrate efficient co-expression and subcellular localization across cell types. Scale bars, 10 µm. **H,** Low-magnification overview of rat brain organotypic coronal slice (300 µm thick, cultured 1 week) transfected with mRNA cocktail encoding LAMP1-HaloTag, mScarlet3-Sec61β, and 4xCOX8-mStayGold. Robust fluorescent labeling is evident throughout the slice. Scale bar, 200 µm. **I,** High-magnification view of (**H**) showing subcellular resolution of lysosomes (magenta), ER (red), and mitochondria (green) within individual neurites in intact tissue architecture. Scale bar, 10 µm.

To quantify co-transfection efficiency, we used the nuclear mStayGold signal to segment individual nuclei and measured the proportion that also expressed mScarlet3, HaloTag, or mTagBFP2 as different mRNA combinations were mixed in equal amounts (**Fig. 5C** and **Supplementary Fig. 3B**). As a control, cells transfected with nuclear mStayGold mRNA alone were used to establish a fluorescence intensity threshold based on the resulting signal distribution. No expression of mScarlet3, HaloTag, or mTagBFP2 was detected in these controls, confirming the specificity of the quantification method (**Fig. 5C, A only**).

When nuclear mStayGold (**A**) and nuclear mScarlet3 (**B**) mRNAs were co-delivered, approximately 94% of mStayGold-positive nuclei also expressed mScarlet3 (**Fig. 5C, A+B**). With three constructs (mStayGold (**A**), mScarlet3 (**B**), and HaloTag (**C**)) about 99% of mStayGold-positive nuclei expressed mScarlet3, and 91% expessed HaloTag, corresponding to roughly 90% co-transfection efficiency across the three mRNAs. Finally, when four constructs (mStayGold (**A**), mScarlet3 (**B**), HaloTag (**C**), and mTagBFP2 (**D**)) were co-transfected at equal ratio, 100% of mStayGold-positive nuclei also expressed mScarlet3, 96% expressed HaloTag, and 93% expressed mTagBFP2. Notably, despite equal mStayGold mRNA input across all conditions, its nuclear fluorescence intensity progressively decreased as more distinct mRNAs were co-delivered, suggesting that total mRNA uptake or translation per cell may be limited by a delivery or expression threshold **(Supplementary Fig.**LJ**3C**).

Having established that multiple transcripts can be effectively co-delivered and co-expressed within single cells, we next applied this strategy to simultaneously label distinct subcellular compartments. mRNAs incorporating the m5C modification and a CleanCap M6 structure were designed to encode LAMP1-HaloTag for lysosomes, mScarlet3-Sec61β for the endoplasmic reticulum, 4xCOX8-mStayGold for mitochondria, and mTagBFP2-SKL for peroxisomes. When transfected into diverse primary cell types, including adult mouse hepatocytes, rat hippocampal neurons, human skeletal muscle cells, human monocyte-derived macrophages, and mouse primary myotube, the mRNA cocktail yielded robust co-expression with high efficiency (**Fig. 5D-G** and **Supplementary Fig. 3D**). This enabled simultaneous live-cell visualization of organelle dynamics and inter-organelle interactions, extending capabilities of existing multi-organelle imaging strategies^23^. A complete list of the plasmids encoding organelle-targeted fluorophores, structural markers, and biosensors (the *4DCP*lJ*mRNAbow*lJ*Collection*) is provided in **Supplementary**LJ**Table**LJ1.

We next evaluated delivery efficiency in tissue context. To test this, we used rat brain coronal slices (300 µm thick, cultured for 1 week), which were transfected with 2 µg each of LAMP1-HaloTag, mScarlet3-Sec61β, and 4xCOX8-mStayGold mRNAs and imaged the next day. Robust fluorescence was detected throughout the slices (**Fig.**LJ**5H**), and higher-magnification imaging revealed clear subcellular localization of labeled organelles within intact tissue architecture, including prominent labeling in neurites (**Fig.**LJ**5I**).

Together, these results demonstrate that synthetic mRNA cocktails enable multiplexed labeling of subcellular structures, supporting dynamic visualization of organelles and their contacts in both primary cell cultures and complex tissue environments.

### *mRNAbow* enables low toxicity, tunable, and multiplexed mRNA expression in zebrafish embryos

We reasoned that optimizing mRNA design for low toxicity and high stability could extend its utility to whole-organism applications. Microinjection of *in vitro* transcribed mRNA is widely used in aquatic model systems to drive transient gene expression or to generate transgenic lines^24,25^ (**Fig. 6A**). However, toxicity has historically constrained the amount of RNA that can be delivered into embryos^26^. Because the total “tolerable” mRNA load per embryo is constrained, mRNA stability also becomes a limiting factor, particularly for later developmental stages, as injected transcripts progressively degrade. For example, in zebrafish embryos, prior studies have shown that the short lifetime of injected mRNA restricts detectable expression to about 3 days post fertilization (3 dpf) when introduced at the one-cell stage^27^. Based on our findings in mammalian cells, we asked whether the *mRNAbow* approach could support efficient and sustained expression *in vivo*.

**Figure 6.**
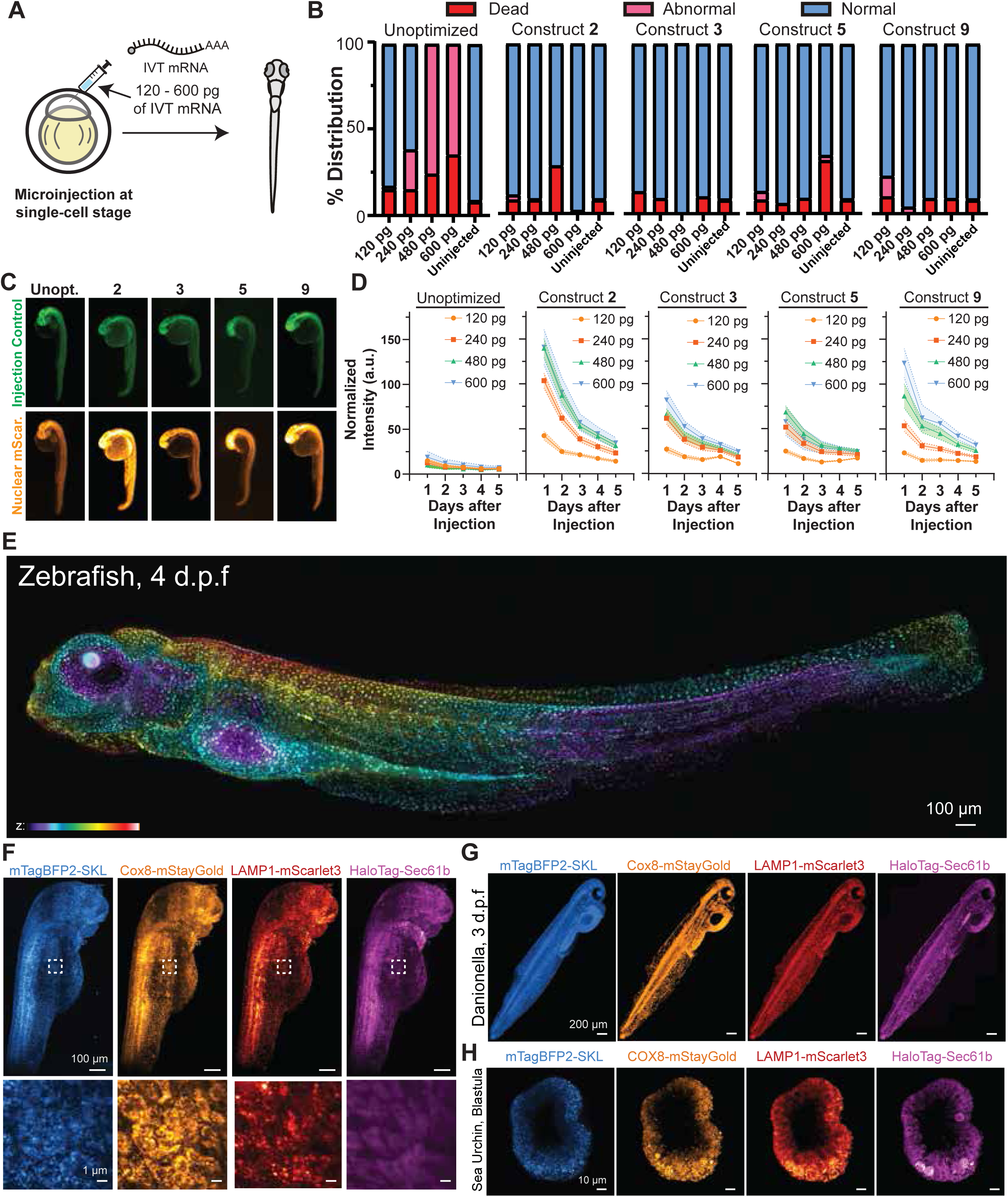
*mRNAbow* enables low toxicity, tunable, and multiplexed mRNA expression in embryos. A, Schematic illustrating microinjection of IVT mRNAs into single-cell zebrafish embryos. **B**, Toxicity assessment of zebrafish embryos injected with increasing amounts of IVT mRNAs by showing % distribution of dead (red), abnormal (magenta), and normal (blue). Embryos were injected with unoptimized mRNA (120 pg, *n* = 39; 240 pg, *n* = 52; 480 pg, *n* = 49; 600 pg, *n* = 51), construct **2** (CleanCap M6, unmodified; 120 pg, *n* = 60; 240 pg, *n* = 75; 480 pg, *n* = 14; 600 pg, *n* = 40), construct **3** (CleanCap M6 + m1Ψ; 120 pg, *n* = 23; 240 pg, *n* = 32; 480 pg, *n* = 49; 600 pg, *n* = 51), construct **5** (CleanCap M6 + m5C; 120 pg, *n* = 35; 240 pg, *n* = 43; 480 pg, *n* = 21; 600 pg, *n* = 32), or construct 9 (CleanCap M6 + 5moU; 120 pg, *n* = 47; 240 pg, *n* = 52; 480 pg, *n* = 32; 600 pg, *n* = 22). Uninjected embryos served as controls (*n* = 247). **C**, Representative images of zebrafish embryos at 1 dpf injected with an internal injection control (CleanCap M6 + m5C nuclear mStayGold; green, top) and nuclear mScarlet3 mRNA variants (bottom; from left to right: unoptimized, construct **2**, construct **3**, construct **5**, construct **9**). **D**, Normalized nuclear mScarlet3 fluorescence intensity relative to mStayGold signal following injection of 120 pg (yellow), 240 pg (red), 480 pg (green), or 600 pg (blue) mRNA, measured over 5 days post fertilization for unoptimized and modified constructs (**2**, **3**, **5**, and **9**). **E**, Color-coded Z-projection of a zebrafish larva at 4 dpf expressing nuclear mScarlet3. Scale bar, 100 µm. **F**, Maximum intensity projection of a zebrafish embryo at 2 dpf expressing mTagBFP2-SKL (blue), 4xCOX8-mStayGold (orange), LAMP1-mScarlet3 (red), and HaloTag-Sec61β labeled with JF635-HTL (magenta). Right, higher-magnification view of the boxed region highlighting subcellular organelle structures. Scale bars, 100 µm (overview) and 1 µm (zoom). **G**, *Danionella cerebrum* larva at 3 dpf expressing mTagBFP2-SKL (blue), 4XCOX8-mStayGold (orange), LAMP1-mScarlet3 (red), and HaloTag-Sec61β labeled with JF635-HTL (magenta). Scale bar, 200 µm. **H**, Fixed *Strongylocentrotus purpuratus* blastula at 24 hours post fertilization expressing mTagBFP2-SKL (blue), 4xCOX8-mStayGold (green), LAMP1-mScarlet3 (yellow), and HaloTag-Sec61β labeled with JF635-HTL (magenta). Scale bar, 10 µm.

To first evaluate toxicity, we injected zebrafish embryos at the single cell stage with mRNA encoding nuclear mScarlet3. Transcripts generated by a standard *in vivo* transcription protocol, using unmodified nucleotides, a non-engineered T7 polymerase, and unmodified primers, were co-transcriptionally capped with ARCA^9^. Embryos microinjected with increasing doses of this transcript showed clear toxicity past 120 pg (**Fig. 6B**, Unoptimized), consistent with previous reports^26^. In contrast, mRNA produced with the *mRNAbow* workflow, which utilizes an engineered T7 polymerase, 2’O-methylated primers, and CleanCap M6, exhibited no detectable toxicity at the same dose, even in the absence of nucleotide modifications (**Fig. 6B**).

We next compared translation efficiency across a set of modified and unmodified mRNAs. To normalize embryo to embryo variability in microinjection, each version of mScarlet3 mRNA was co-injected with a 120 pg of m5C-modified nuclear mStayGold mRNA, which served as an internal normalization reference. Normalized fluorescence was quantified at 1 day post fertilization (1Ldpf). As expected, uncapped transcripts produced no detectable fluorescence, confirming that zebrafish embryos also require a functional cap structure for translation initiation (**Supplementary Fig. 4A**).

Unexpectedly, nucleotide modifications that strongly improve expression in post-mitotic and terminally differentiated mammalian cells, including m1Ψ and m5C, yielded weaker fluorescence compared to mRNA lacking nucleotide modifications although all optimized transcripts performed better than control mRNA generated using standard *in*lJ*vitro* transcription protocols described above **(Fig.**LJ**6C,**LJ**D)**. Among all variants, unmodified mRNA produced the strongest mScarlet3Lsignal, followed byL5moU, suggesting that early zebrafish embryos regulate RNA differently from differentiated mammalian cells.

We then examined how expression changed over developmental time. Although unmodified mRNA drove strong initial fluorescence, the signal diminished rapidly after 2 dpf (**Fig. 6D**). By contrast, 5moU modified mRNA maintained expression longer, showing a gradual decrease after 2dpf and uniform distribution pattern through 4Ldpf (**Fig. 6D and 6E**). These results indicate that specific nucleotide modifications influence expression stability over embryo development, providing an opportunity to tailor nucleotide modifications to experimental timelines and the desired stability of the encoded product.

As a functional test of sustained expression over development, we microinjected 400 pg of 5moU-modified, M6-capped GCaMP7f mRNA into zebrafish embryos at the one-cell stage. GCaMP7f has previously been shown to function robustly in zebrafish^28^. At 4 dpf, larvae were anesthetized and calcium dynamics were monitored by imaging GCaMP7f fluorescence. Expression remained readily detectable at this stage, though fluorescence intensity varied across tissues, potentially reflecting spatial differences in calcium activity **(Video**LJ**S8)**. Spontaneous calcium transients were observed in multiple regions, including the skin and heart **(Video**LJ**S8**), demonstrating that optimized mRNAs can sustain functional reporter expression over developmental timescales suitable for *in vivo* imaging.

Finally, we tested whether *mRNAbow* supports multiplexed expression *in vivo* in zebrafish embryos. A mixture of four 5moU modified mRNAs encoding mTagBFP2-SKL, 4xCox8-mStayGold, LAMP1-mScarlet3, and HaloTag-Sec61β was microinjected (150 pg each) at the single cell stage in the zebrafish embryo. Distinct fluorescence patterns marking peroxisomes, mitochondria, lysosomes, and endoplasmic reticulum were readily detected in a 2 dpf embryo (**Fig. 6F** and **Supplementary Fig. 4B**). These results show that *mRNAbow* enables simultaneous, non-toxic expression of multiple reporters, allowing visualization of several organelles within the same embryo.

### *mRNAbow* enables multiplexed expression in embryos with limited genetic accessibility

We next explored whether the *mRNAbow* approach could be extended to species with limited genetic tools. *Danionella cerebrum* is an emerging teleost model organism notable for its lifelong optical transparency, compact brain, and close evolutionary relationship to zebrafish^29^, making it well suited for *in vivo* imaging and neural circuit analysis. However, the development of stable transgenic lines in *Danionella* remains limited. Microinjection of stable IVT mRNAs, therefore, offers a complementary approach in this system by enabling fast, transient, and combinatorial expression without the need for germline modification. To test whether our *mRNAbow* strategy is compatible with *Danionella*, we microinjected a mixture of four 5moU modified mRNAs encoding mTagBFP2-SKL, 4xCOX8-mStayGold, LAMP1-mScarlet3, and HaloTag-Sec61β at the single-cell stage and performed imaging at 3 days post fertilization. The larvae exhibited robust and specific fluorescent labeling across multiple subcellular compartments, demonstrating efficient expression and compatibility of *mRNAbow* in *D. cerebrum* (**Fig. 6G** and **Supplementary Figure 5**).

Because many organelle targeting sequences are evolutionarily conserved, we next asked whether this approach could be extended to more distantly related basal deuterostomes. To test this, we microinjected the same set of IVT mRNAs (∼2 pL of 150 pg each) into *Strongylocentrotus purpuratus* (purple sea urchin) zygotes. Fluorescent signal was detectable beginning 4 hours post fertilization and persisted through larval stages at 3 days post fertilization, with clear labeling of distinct subcellular compartments in fixed embryos (**Fig. 6H** and **Supplementary Figure 6, 7,** and **8**).

Together, these results demonstrate that *mRNAbow* enables efficient, multiplexed organelle labeling across evolutionarily diverse species and can be readily applied beyond established genetic model systems.

## Discussion

In this study, we present *mRNAbow*, a framework for the design, production, and application of *in vitro* transcribed (IVT) mRNAs as a versatile toolkit for cell biology. By systematically evaluating capping strategies and nucleotide modifications across diverse systems, we establish design principles that enable robust protein expression in contexts where plasmid- or virus-based approaches are limited. Our results reaffirm that a 5′ cap is essential for translation, as capped mRNAs consistently produced strong protein expression, whereas uncapped transcripts failed across all systems tested. In terminally differentiated and post-mitotic mammalian cells, 5-methylcytidine (m5C) emerged as a broadly compatible modification, supporting robust expression across multiple primary cell types without detectable cytotoxicity or innate immune activation. In contrast, experiments involving microinjection into single-cell zebrafish embryos revealed that unmodified and 5-methoxyuridine (5moU)-modified mRNAs outperformed m5C-modified transcripts. Together, these findings establish capping as a universal requirement for IVT mRNA function, identify m5C as a practical modification for applications in differentiated mammalian cells, and highlight 5moU as an effective nucleotide modification for single-cell embryo injection. Collectively, they underscore the potential of IVT mRNAs as plug-and-play tools for multiplexed experimental applications.

The necessity of m7G cap is well supported by decades of work identifying the m7G cap as the molecular “gatekeeper” of cytoplasmic translation^30,31^. In our hands, CleanCap-derived Cap1 RNAs consistently outperformed uncapped controls. Recent innovations introducing N6-methyladenosine (m6A) at the +1 position provide a mechanistic basis for the observed sustained signals, suggesting that m6A at this site may impede decapping and prolongs RNA stability as previously suggested^17^. This enhanced stability, combined with optimized nucleotide incorporation, yielded transcripts that balance durability with broad compatibility, offering a practical solution for diverse biological systems.

Our template design and transcription strategy also contributed to the absence of stress pathway activation. Promoter-independent initiation by T7 polymerase is a known source of double-stranded RNA contaminants^32,33^. We minimized this by combining engineered T7 variants with 2’-O-methylated DNA template ends, reducing dsRNA byproducts without sacrificing yield. Consistent with previous reports^19^, we observed no evidence of stress responses such as phosphorylation of eIF2α, RIG-I activation, or IRF3 phosphorylation, even with unmodified capped RNAs after 24-hr post-transfection (**Fig. 1E**). The impact of the *in vitro* transcription workflow was further underscored by the toxicity patterns observed in zebrafish embryos. mRNA generated using a conventional *in vitro* transcription workflow, incorporating unmodified nucleotides, a nonengineered T7 polymerase, and template generated from primers lacking 2′-O-methylation, exhibited clear dose-dependent toxicity with adverse effects apparent above 120 pg, consistent with previous reports^26^. In contrast, mRNA produced using the *mRNAbow* workflow, which integrates an engineered T7 polymerase, template generated from 2′-O-methylated primers, and CleanCap M6, showed no detectable toxicity at the same dose, even in the absence of nucleoside modifications. This contrast suggests that dsRNA contaminants or transcriptional byproducts, rather than the absence of modified nucleotides, are key drivers of toxicity in embryo systems.

Interestingly, capped and uncapped RNAs displayed similar intracellular delivery kinetics measured by RT-qPCR, with both reaching a plateau in abundance after approximately 8 hours and remaining detectable for over 24 hours (**Fig. 3A**). The divergence in protein output (**Fig. 2C**) therefore implicates mRNA modification as a determinant of translational efficiency rather than mRNA stability in mammalian cells. Prior work has shown that N1-methylpseudouridine (m1Ψ) can enhance decoding efficiency in some contexts but also increase translational frameshifting^16^. Consistent with this, in m1Ψ-containg construct (**3**), we occasionally observed mislocalized cytosolic fluorescence of nuclear mScarlet3, suggesting that a fraction of the translated product may be mislocalized due to frameshift-induced disruption of targeting sequences. These observations highlight that the choice of mRNA modification must be carefully considered to maintain both efficient and accurate protein expression. Across all conditions, m5C consistently yielded robust and correctly localized expression in hepatocytes, neurons, macrophages, muscles, and organoid, supporting previous findings that m5C does not induce translational frameshifts ^16^.

A methodological highlight of *mRNAbow* lies in showing that IVT mRNAs can serve as modular building blocks for multiplexed experiments. Capitalizing on the capacity of lipid formulations to co-deliver multiple transcripts^22^, we achieved rapid, simultaneous expression of organelle markers and biosensors in primary cells and tissue slices. This approach contrasts with viral delivery systems, which are limited by packaging constraints and slow expression kinetics. With *mRNAbow*, new marker combinations can be tested within hours, greatly accelerating exploratory imaging studies and facilitating experiments in human samples where biosafety and time are limiting factors.

The same principle holds true *in vivo*, where multiplexed mRNA microinjection offers distinct advantages over traditional genetic approaches, even in well-established model systems like zebrafish. Injection of higher doses of optimized mRNAs substantially improve signal to noise in live imaging experiments, especially during early development. Moreover, increased transcript longevity enabled fluorescence detection well beyond the typical 3 dpf limit of conventional mRNAs. This approach allows multiplexed organelle imaging that would otherwise require multiple transgenic lines generated over months or years. These benefits are particularly transformative for emerging research organisms lacking robust genetic tools or with life history constraints that preclude multigenerational breeding. The evolutionary conservation of organelle targeting domains further facilitates direct transfer of reporter designs across species.

By pairing a standardized design framework with an open-access library of reporters, structural markers, and biosensors (“the *4DCP mRNAbow Collection*”), we aim to lower the barrier for adoption of mRNA tools in cell and developmental biology. Making these reagents and protocols broadly available promotes reproducibility and accessibility, enabling research groups without prior mRNA expertise to readily deploy this system. Future directions include the exploration of additional nucleotide modifications, improved delivery chemistries, and broader organismal applications. Optimization of lipid- or polymer-based delivery vectors, particularly those incorporating tissue-targeting capabilities, could further expand the reach of IVT mRNA technologies.

Looking forward, *mRNAbow* offers a foundation not only for multiplexed imaging but also for the transient delivery of functional proteins, including genome editing enzymes, optogenetic actuators, and transcriptional regulators. By bridging advances in therapeutic mRNA engineering with the needs of experimental biology, this framework provides a versatile, accessible, and rapidly adaptable platform to interrogate the spatial and temporal organization of living systems.

## Acknowledgments

We thank Lippincott-Schwartz lab members, Light microscopy core, Janelia Shared Resources, Janelia Aquatics, Molecular Genomics, Daniel Feliciano, Alejandro Aguilera Casterejon, Raghabendra Adhikari, Jeremy Hasseman, Triveni Menon, Tanishqua Duarah, Virginia Ruetten, Kelsey Voge, and others for scientific discussions and helpful resources. We also thank Luke Lavis and the Open Chemistry team for providing the JF dyes used in our experiments. This work was supported by the Howard Hughes Medical Institute.

## Author Contributions

Conceptualization, H.C. and J.L.-S.; methodology, H.C.; investigation, H.C., C.H., W.K., M.D.T., P.N., D.W., A.T., S.R.-E., H.W., D.C., E.Y.S. C.S. D.Q.M.; resources, W.K., I.L.W., A.T., I.E.M, J.L.S., D.M., J.L-S; data curation, H.C., C.H, W.K; software, H.C. and C.H; formal analysis, H.C. and C.H; supervision, H.C and J.L.-S.; writing, H.C.; manuscript review and editing, all authors

## Data and materials availability

All plasmids will be made available through Addgene, but any early requests can be made to the corresponding authors.

**Video S1. Time-lapse imaging of nuclear mScarlet3 expression.** Live-cell fluorescence microscopy showing progressive nuclear accumulation of mScarlet3 over time following mRNA transfection over 16 hours. Each frame was acquired at 10 min intervals. The scale bar indicates 200 µm.

**Video S2. Time-lapse imaging of mScarlet3-Sec61**β **expression.** Live-cell fluorescence microscopy showing progressive ER accumulation of mScarlet3 over time following mRNA transfection over 12 hours. Each frame was acquired at 10 min intervals. The scale bar indicates 100 µm.

**Video S3. Tracking of EB3 dynamics in a cultured rat hippocampal neuron.** Time-lapse microscopy visualizing EB3 comets along neuronal microtubules. Each frame was acquired at 2.4 sec intervals. The scale bar indicates 10 µm.

**Video S4. Tracking of EB3 dynamics in a cultured rat hippocampal neuron.** Time-lapse microscopy visualizing EB3 comets along neuronal microtubules. Trajectories highlight the orientation and speed of microtubule plus-end growth within axons and dendrites. Each frame was acquired at 2.4 sec intervals. The scale bar indicates 10 µm.

**Video S5. Tracking of EB3 dynamics in a human skeletal muscle cell.** Time-lapse microscopy visualizing EB3 comets along microtubules in muscles. Each frame was acquired at 4.8 sec intervals. The scale bar indicates 15 µm.

**Video S6. GCaMP8s calcium dynamics in human iCardiomyocytes.** Live imaging of calcium transients in induced pluripotent stem cell-derived (iPSC) cardiomyocytes expressing GCaMP8s. Fluorescence fluctuations correspond to rhythmic calcium oscillations during spontaneous contractions. Each frame was acquired at 0.93 sec intervals. The scale bar indicates 10 µm.

**Video S7. GCaMP8s calcium dynamics in rat brain slice.** Live imaging of calcium transients in rat brain slice expressing GCaMP8s. Fluorescence fluctuations correspond to rhythmic calcium oscillations during spontaneous contractions. Each frame was acquired at 2.5 sec intervals. The scale bar indicates 100 µm.

**Video S8. GCaMP7f in Zebrafish larvae.** Live calcium imaging of zebrafish larvae at 4 days post fertilization. Color-coded Z projections are shown with depth encoded from 0 to 205 µm. Scale bar, 100 µm. Time between frames is 26 seconds.

**Supplementary Fig 1. Uncropped Western Blots of Fig. 1E**.

**Supplementary Fig. 2 Cell type–specific preferences for nucleotide modifications in mRNA constructs.** Expression of nuclear mScarlet3 expression in **A,** Adult mouse hepatocyte. Scale bars, 250 µm, with DNA stain **B**, Human Skeletal Muscle. Scale bars, 50 µm, with DNA stain **c**, rat hippocampal neurons. Scale bars, 50 µm, with neurofilament M antibody staining **d**, iPSC-derived Macrophage. Scale bars, 50 µm, with DNA stain. **C**, iPSC-derived Brain organoids. Scale bars, 250 µm, with DNA stain. **1** = no modification, no cap, **2** = no modification, M6 Cap, 3 = N1-methylpseudouridine (m1Ψ), M6 Cap, **5** = 5-methylcytidine (m5C), M6 Cap, **7** = N1-methylpseudouridine (m1Ψ) + 5-methylcytidine (m5C), M6 Cap **9** = 5-methoxy-uridine (5moU), M6 Cap. Scale bars, 50 µm.

**Supplementary Fig. 3 Multiplexed mRNA delivery and reporter co-expression in skeletal muscle, U-2 OS cells, and primary myotubes. A,** Skeletal muscle expressing EB3-mStayGold. Scale bar, 10 µm. **A–C**, Images of nuclei (**B**) and intensity quantification (**C**) from mStayGold-segmented nuclei (**A**) and co-expression of mScarlet3 (**B**), HaloTag (**C**), and mTagBFP2 (**D**) under one- (**A only**), two- (**A+B**), three- (**A+B+C**), and four-mRNA (**A+B+C+D**) delivery conditions in U-2 OS cells. Scale bars, 50 µm. **D**, Mouse primary myotube derived from ROSA^mT/mG^ expressing HaloTag-Sec61β, tdTomato (from ROSA^mTmG^), LAMP1-mStayGold, and mTagBFP2-SKL.

**Supplementary Fig. 4 Validation of mRNA expression and four-color organelle imaging in zebrafish embryos. A.** Images of zebrafish embryos at 1 day post fertilization (dpf) injected with 120 pg or 240 pg of uncapped, unmodified nuclear mScarlet3 mRNA. **B.** Zoomed-in images of a 2 dpf zebrafish injected with 150 pg of mTagBFP2-SKL (cyan), 4xCOX8-mStayGold (yellow), LAMP1-mScarlet3 (red), and HaloTag-Sec61β labeled with 5 µM JF635-HTL (magenta). Scale bar, 50 µm.

**Supplementary Fig 5. Four-color organelle imaging in *Danionella cerebrum*.** High-magnification view of a 3 dpf *D. cerebrum* larva injected with 150 pg of mTagBFP2-SKL (cyan), 4xCOX8-mStayGold (yellow), LAMP1-mScarlet3 (red), and HaloTag-Sec61β labeled with 5 µM JF635-HTL (magenta). Scale bar, 50 µm.

**Supplementary Fig. 6. Purple Sea Urchin Blastula. A** Fixed *Strongylocentrotus purpuratus* embryo at the blastula (24 hours post fertilization) injected with equivalent amount of Firefly mRNA do not show fluorescence. Scale bars, 10 µm. **B.** Fixed *Strongylocentrotus purpuratus* embryo at the blastula (24 hours post fertilization) expressing mTagBFP2-SKL (blue), 4xCOX8-mStayGold (green), LAMP1-mScarlet3 (yellow), and HaloTag-Sec61β labeled with JF646-HTL (magenta). Scale bars, 50 µm. Images are shown from top to bottom as sequential Z planes.

**Supplementary Fig. 7. Purple Sea Urchin Gastrula.** Fixed *Strongylocentrotus purpuratus* embryos at gastrula (48 hours post fertilization) stage expressing mTagBFP2-SKL (blue), 4xCOX8-mStayGold (green), LAMP1-mScarlet3 (yellow), and HaloTag-Sec61β labeled with JF646-HTL (magenta). Vegetal view is presented. Scale bars, 50 µm. Images are shown from top to bottom as sequential Z planes.

**Supplementary Fig. 8. Purple Sea Urchin Larvae.** Fixed *Strongylocentrotus purpuratus* larva (72 hours post fertilization) expressing mTagBFP2-SKL (blue), 4xCOX8-mStayGold (green), LAMP1-mScarlet3 (yellow), and HaloTag-Sec61β labeled with JF646-HTL (magenta). Scale bars, 50 µm. Images are shown from top to bottom as sequential Z planes.

## Materials and Methods

Animal work was conducted according to the Institutional Animal Care and Use Committee guidelines of Janelia Research Campus (IACUC protocol #25-0268, #24-0258, # 24-0261.06).

### Reagents

CleanCap M6 was purchased from TriLink. N1-methyl ΨTP and 5m-CTP was purchased from TriLink. 5-mo UTP was purchased from NEB. ATP, GTP, CTP, and UTP solutions were purchased from NEB. Superase-In RNase inhibitor was purchased from Thermo Fisher Scientific. RNase free DNase I was purchased from NEB. Ethanol was purchased from Sigma Aldrich. Hoechst 33342 was purchased from Thermo Fisher.

### DNA Template Preparation

Template DNA for in vitro transcription was amplified by PCR using primers 5’-mGmC(T)_44_-3’ and 5’-CGAAATTAATACGACTCACTATAAGGAATAA-3’ with Q5 Hot Start High-Fidelity Master Mix (NEB) and 1 ng of plasmid template and 500 nM of primers. Reactions were performed with an annealing temperature of 64 °C. PCR products were verified by agarose gel electrophoresis, purified using the Monarch PCR & DNA Cleanup Kit (NEB), and eluted in RNase-free water. DNA concentration was determined by NanoDrop spectrophotometry, and purified products were used directly for *in vitro* transcription.

### In vitro transcription

*In vitro* transcription (IVT) was carried out using the Takara PrimeCap T7 RNA polymerase kit, which employs a mutated T7 RNA polymerase engineered to minimize double-stranded RNA (dsRNA) by-products, thereby improving transcript purity and translation efficiency. Co-transcriptional capping was achieved with CleanCap M6 cap analogs (TriLink). For chemical nucleotide modifications, cytidine triphosphate (CTP) was substituted with 5-methyl-CTP, and uridine triphosphate (UTP) was replaced with either N1-methylpseudouridine triphosphate (N1-methyl-ΨTP) or 5-methoxyuridine triphosphate (5mo-UTP). Reactions were assembled with ≥500 ng of DNA template and supplemented with SUPERase•In RNase inhibitor (1 U/µL). IVT was conducted at 37 °C for 2 h, followed by treatment with 4 µL RNase-free DNase I at 37 °C for 15 min to remove template DNA. Transcripts were purified using the NEB Monarch RNA Cleanup Kit (NEB) and eluted in RNase-free water. RNA integrity and purity was verified by agarose gel electrophoresis, and samples were diluted to the desired concentration, aliquoted into RNase-free tubes, and stored at −80 °C until use. For a detailed protocol, see the **Supplementary Information**.

Unoptimized mRNA was synthesized using the mMESSAGE mMACHINE™ T7 Transcription Kit (Thermo Fisher Scientific, AM1344). *In vitro* transcription reactions were incubated at 37 °C for 4 hours, followed by DNase treatment with 1 µL TURBO DNase (2 U/µL) for an additional 15 minutes at 37 °C to remove template DNA. Transcribed mRNA was purified using the MEGAclear™ Transcription Clean-Up Kit (Invitrogen, AM1908) according to the manufacturer’s instructions.

### mRNA Transfection

*In vitro*-transcribed mRNAs were transfected using Lipofectamine MessengerMAX (ThermoFisher) at a 3:1 reagent-to-RNA ratio. Lipofectamine MessengerMAX was first diluted in Opti-MEM (ThermoFisher) pre-equilibrated to room temperature and incubated for 10 min. In parallel, IVT mRNA was diluted in an equal volume of room temperature Opti-MEM. The two solutions were then combined, mixed gently by pipetting, and incubated for 5 min at room temperature to allow complex formation. The transfection mixture was added directly to the culture medium or brain slice preparation, and cultures were gently agitated to ensure even distribution. For multi-construct transfection, mRNAs encoding different proteins were pre-mixed in Opti-MEM (ThermoFisher) and then combined with the transfection reagents using the same ratio of mRNA:Lipofectamine as described above.

### qPCR protocol

Quantitative reverse transcription PCR (RT-qPCR) was conducted using the Luna Cell Ready One-Step RT-qPCR Kit (New England Biolabs, E3030) following the manufacturer’s instructions. U-2 OS cells were seeded in 96-well plates and transfected with 100 ng of the respective mRNAs using a 1:3 ratio of mRNA to Lipofectamine MessengerMax reagent in 10 µL of Opti-MEM medium. Cells were harvested at 4-, 8-, 16-, and 24-hours post-transfection. At each time point, cells were lysed with the Luna Cell Ready Lysis Module to obtain RNA-containing lysates without further purification. Lysates were briefly centrifuged and used directly as templates for one-step RT-qPCR.

Each reaction contained cell lysate, Luna Cell Ready Reaction Mix (NEB), and gene-specific primers targeting the transcripts of interest as indicated in **Supplementary Table 2**. *ACTB* expression was used as an internal reference for normalization. Reactions were performed in technical triplicate using a Roche real-time PCR system. Quantification cycle (Cq) values were automatically determined by the instrument software. Relative mRNA expression levels were determined using the ΔΔCq method by comparing each sample to the no-transfection control and ACTB. qPCR was performed with two different sets of primers specific to nuclear mScarlet3, and expression levels were normalized to the highest value for each mRNA across all time points.

### U-2 OS Cell culture

U-2 OS (ATCC) cells were cultured in phenol-red free DMEM supplemented with 10% fetal bovine serum (Corning), 2 mM L-glutamine (Corning), and 100 IU penicillin and 100 µg/mL streptomycin (CellGro) at 37°C and 5% CO_2_. For imaging and transfection experiments, an 18-well coverslip-bottom imaging chamber (CellVis, C18-1.5H) was coated with Matrigel (Corning, 356231) diluted 1:10 in Opti-MEM (ThermoFisher, 11058021). U-2 OS cells were seeded at approximately 15,000 cells per well one day before transfection. To quantify transfection efficiency, images were acquired using a Zeiss LSM 980 microscope equipped with a Plan-Apochromat 20x/0.80 M27 objective.

### Rat Hippocampal Neuronal Culture

Dissociated hippocampal neurons were isolated from P0 Sprague-Dawley rat pups (Charles River). Hippocampi were dissected and enzymatically digested with papain in dissection buffer (10 mM HEPES in Hanks’ balanced salt solution, HBSS). Following digestion, the tissue was gently triturated in minimum essential medium (MEM) supplemented with 10% fetal bovine serum (FBS) and passed through a 40 μm cell strainer. Cell density and viability were assessed by trypan blue exclusion using a Vi-Cell Blu cell counter. To prepare the suspension, plating medium (DMEM with 10% FBS) was mixed with NbActiv4 (Transnetyx) at a 1:10 ratio, diluting the cells to a final concentration of 100,000 cells/mL. Approximately 50,000-75,000 neurons were seeded onto poly-D-lysine-coated coverslips in 24-well plates. Cultures were maintained by replacing half of the medium with fresh NbActiv4 once per week until the neurons were used. To quantify transfection efficiency, images were acquired using a Zeiss LSM 980 microscope equipped with a Plan-Apochromat 20x/0.80 M27 objective. Cells were fixed with 4% paraformaldehyde (PFA) and 4% sucrose in PBS for 15 minutes, followed by three washes with PBS. Permeabilization was performed using 0.1% Triton X-100 in PBST for 15 minutes. The samples were then blocked with SuperBlock Blocking Buffer in PBS (ThermoFisher) for 1 hour.

After blocking, cells were incubated overnight at 4°C with the primary antibody against neurofilament M (NF-M, Encore, RPCA-NF-M). The samples were rinsed three times for 5 minutes each with PBST, then incubated for 1 hour with the goat anti-rabbit IgG (H+L) Alexa Fluor Plus 488 (ThermoFisher). Finally, the cells were washed three times with PBST before imaging. For higher-resolution images, we used Zeiss LSM980 with a C-Apochromat 63x/1.2 W Corr M27 objective.

### Adult mouse hepatocyte culture

Primary hepatocytes were isolated from 7-8-week-old C57BL/6 mice (Charles River Laboratories) using a two-step collagenase perfusion method via vena cava cannulation, as previously described^34^, with minor modifications. Briefly, 20 mL of enzyme digestion buffer (25 mM HEPES) was perfused through the liver to enhance cell yield. Hepatocytes were purified following the published protocol and optimized for subsequent culture conditions. 24-well plates were coated with 250 µL of 50 µg/mL collagen (Sigma-Aldrich, Cat. #122-20) for 2 h at 37 °C, washed three times with PBS, and air-dried. Cells were seeded at a density of 8 x 10L cells per well in 500 µL of cell attachment medium and incubated at 37 °C with 10% CO_2_ + 16% O_2_ for 3 h. The medium was then replaced with 500 µL of hepatocyte maintenance medium, which was refreshed daily. Cultures were transfected with 250 ng of mRNAs with Lipofectamine MessengerMax reagent diluted at 1:3 ratio 24 h post-seeding. To quantify transfection efficiency, we used EVOS M5000 system equipped with a 20x objective. For higher-resolution images, we used Zeiss LSM980 with a C-Apochromat 63x/1.2 W Corr M27 objective.

### Human skeletal muscle culture

Primary normal human skeletal myoblasts (HSkM; ThermoFisher, A12555) were thawed and differentiated according to the manufacturer’s instructions. Briefly, an 18-well coverslip-bottom imaging chamber (CellVis, C18-1.5H) was coated with Matrigel (Corning, 356231) diluted 1:10 in Opti-MEM (ThermoFisher, 11058021). Frozen vials of HSkM were rapidly thawed in a 37 °C water bath with gentle agitation, then immediately transferred to a sterile 50-mL conical tube containing 10 mL of differentiation medium (D-MEM Basal Medium (Cat. No. 11885-084) supplemented with 2% Horse Serum (Cat. No. 16050-130)). The cryovial was rinsed once with 1 mL of differentiation medium, and the rinse was combined with the cells. The suspension was centrifuged at 180 x g for 5 min at room temperature, the supernatant was aspirated, and the pellet resuspended in 5 mL of fresh differentiation medium by gentle pipetting. Cells were seeded at a density of 48,000 cells per well. Cultures were maintained in a humidified incubator at 37 °C with 5% CO₂, fed every other day with differentiation medium, and allowed to differentiate for 7 days prior to transfection with 100 ng of mRNAs. To quantify transfection efficiency, images were acquired using a Zeiss LSM 980 microscope equipped with a Plan-Apochromat 20x/0.80 M27 objective. For higher-resolution images, we used Zeiss LSM980 with a C-Apochromat 63x/1.2 W Corr M27 objective.

### Primary myoblast isolation and differentiation

Primary myoblasts were isolated from hindlimb and forelimb muscles of adult mice (ROSA^mT/mG^). Dissected muscles were trimmed to remove tendons and fat, minced for 10 min with sterile razor blades, and digested in 0.05% trypsin at 37 °C for 30 min with intermittent mixing. The digestion was quenched with wash medium (DMEM/F12 supplemented with 10% FBS and 1% penicillin-streptomycin (P/S)), and the suspension was centrifuged at 500 x g for 5 min at 4 °C. The pellet was washed twice, then mechanically dissociated using a 5 mL pipette, passed through a 70 µm cell strainer, and centrifuged again under the same conditions.

Cells were resuspended in AmnioMAX II complete medium (Gibco) and seeded onto 10% Matrigel-coated 6 cm dishes at 7x10^5^ cells per well. Cultures were maintained at 37 °C, 5% CO₂, and sub-cultured before reaching 80% confluency. Myoblast enrichment was performed by two rounds of pre-plating: cells were detached with 0.25% trypsin, plated onto uncoated dishes for 1 h, and non-adherent cells were collected and transferred to new T-flasks (passage 1). Cells were cryopreserved at passage 1-2 when sufficient numbers were obtained.

The isolated primary myoblasts were seeded at 1.6x10^4^ cells per well onto 10% Matrigel-coated 18-chamber CellVis plates and cultured for 24 h in proliferation medium. The medium was then replaced with differentiation medium (DMEM/F12 + 2% horse serum + 1% P/S + 1 µM ERK1/2 inhibitor).

### Maintenance of hiPSCs

The KOLF2.1J hiPSC line was used for neural differentiation studies and maintained on vitronectin-coated culture dishes (Corning, Cat. #CLS3535-1EA) in mTeSR+ medium (STEMCELL Technologies). Cells were passaged using ReLeSR (STEMCELL Technologies) according to the manufacturer’s protocol. The WTC-11 hiPSC line was cultured under the same conditions and used for macrophage differentiation experiments.

### Differentiation of cortical organoids

Cortical organoids were generated following the protocol^35^ with minor modifications. Briefly, 4,000 hiPSCs were seeded per well in a round-bottom, ultra-low attachment 96-well plate in stem cell basal medium supplemented with SB431542, dorsomorphin, IBET151, retinoic acid, and CEPT. After 24 h, the medium was replaced with stem cell medium containing SB431542 and dorsomorphin. After an additional 24 h, cultures were switched to IDM supplemented with SB431542 and dorsomorphin. Medium was exchanged every other day until day 10, when organoids were transferred to a ClinoStar2 bioreactor in NMM supplemented with 0.5% Matrigel. On day 20, cultures were switched to A-NMM medium and maintained thereafter. Organoids were used for experiments at day 30 and transfected with 250 ng of mRNAs. To quantify transfection efficiency, images were acquired using a Zeiss LSM 980 microscope equipped with a Plan-Apochromat 20x/0.80 M27 objective.

### iPSC-derived Macrophage differentiation

hiPSCs were first differentiated into monocytes using the STEMdiff Monocyte Kit (STEMCELL Technologies, Cat. #05320). The glass-bottom imaging wells (CellVis, C18-1.5H) were coated with 50 µg/mL poly-D-lysine (PDL) for 1 h at room temperature, rinsed three times with sterile water, and allowed to dry completely. Plates were then incubated with 10 µg/mL fibronectin in DPBS for 1 h at 37 °C, followed by one rinse with PBS. Monocytes were seeded directly into coated imaging wells in ImmunoCult-SF Macrophage Differentiation Medium (STEMCELL Technologies, Cat. #10961) supplemented with 100 ng/mL recombinant human M-CSF (Peprotech, 315-02-10UG). Direct seeding into imaging dishes was critical to maintain optimal cell morphology.

Cultures were maintained at 37°C in a humidified 5% CO₂ incubator. On day 3, half of the medium was replaced with fresh ImmunoCult-SF medium containing 150 ng/mL M-CSF. This replenishment was repeated on day 6. From day 6 to day 13, cells were maintained by replacing media every two days with IMDM supplemented with 10 mM HEPES, 1 mM sodium pyruvate, 5% CTS-SR (ThermoFisher, A2596101), 1x GlutaMAX, 25 ng/mL GM-CSF, and phenol red.

Differentiated macrophages displayed characteristic adherence, spread morphology, and pseudopodia. The transfection was performed with 100 ng of mRNAs. To quantify transfection efficiency, images were acquired using a Zeiss LSM 980 microscope equipped with a Plan-Apochromat10x/0.45 M27 objective.

### Human monocyte isolation and macrophage differentiation

Primary human macrophages were generated from peripheral blood obtained from leukocyte reduction system chambers provided by the Stanford Blood Center. Peripheral blood mononuclear cells (PBMCs) were isolated by Ficoll density gradient centrifugation (Ficoll-Paque PLUS, Cytiva 17144002), and CD14⁺ monocytes were enriched using magnetic bead separation (Miltenyi Biotec, 130-090-879) following the manufacturer’s protocol. Purified monocytes were differentiated into macrophages over 7 days in Iscove’s Modified Dulbecco’s Medium (IMDM; ThermoFisher 12440053) supplemented with 5% heat-inactivated AB human serum (Gemini Bio,100-512-100) + 100 ng/mL GM-CSF (Miltenyi Biotec, 130-095-372). To quantify transfection efficiency, images were acquired using a Zeiss LSM 980 microscope equipped with a Plan-Apochromat 20x/0.80 M27 objective. For higher-resolution images, we used Zeiss LSM980 with a C-Apochromat 63x/1.2 W Corr M27 objective.

### Cardiomyocyte differentiation

Human iPSC lines were reprogrammed with blood cells from health donors under the protocol approved by University of Pittsburgh IRB committee (STUDY21100125). For cardiomyocyte differentiation, iPSCs were cultured to confluency and differentiated into beating cardiomyocytes using a monolayer differentiation protocol as previously described. Briefly, confluent iPSCs were rinsed with DPBS and treated with 6 μM CHIR99021 in RPMI 1640 and B27 supplement minus insulin (RPMI+B27-Insulin) for 48 h. Subsequently, cells were switched back to RPMI+B27-Insulin for 24 h and then exposed to 5 μM IWR with RPMI+B27-Insulin for 48 h. Afterward, cells were placed in RPMI+B27-Insulin for 48 h and finally cultured in RPMI 1640 and B27 supplement for 48 h. The medium was changed every 48 h until beating cells were observed. To prepare the cells for transfection and imaging, iPSC-CMs at D30 after differentiation will be reseeded in Matrigel (Corning) -precoated 18 well CellVis chamber (CellVis, C18-1.5H) at a density of 200 K cells per cm^2^. For time-lapse images, we used Zeiss LSM980 with a C-Apochromat 63x/1.2 W Corr M27 objective.

### Rat brain slice culture

Organotypic hippocampal slice cultures were prepared from postnatal day 4-6 Sprague Dawley rat pups. Animals were decapitated in accordance with institutional animal care and use committee (IACUC) guidelines. Brains were rapidly removed and placed in ice-cold dissection medium consisting of HBSS supplemented with 10 mM glucose and 1% P/S. The brain was isolated and cut into 300 µm coronal sections using a vibratome (Leica VT 1200S). Slices were transferred onto Millicell-CM membrane inserts (Millipore) placed in 35 mm culture plates containing 1 mL slice culture medium (78.5% MEM, 250 μM Ascorbic Acid, 1 mM MgSO_4_, 1 mM CaCl_2_, 15% heat-inactivated horse serum, B27, 25 mM HEPES, and GlutaMAX). Cultures were maintained at 37 °C in a humidified 5% CO₂ incubator, with medium exchanged every 2-3 days. Brain slices were maintained for at least 7 days in vitro (DIV) prior to RNA transfection. For transfection, *in vitro*-transcribed mRNA-lipid complexes were prepared and applied directly onto each slice at a total of 2 µg per mRNA construct in 50 µL. The complexes were delivered dropwise (3 µL per drop) across the surface of the slice to ensure even distribution. After 16-hr post-transfection, the transfected slice was imaged on upright Zeiss 980 equipped with Fluor 5x/ 0.27 M27 air objective and 20x W Plan-Apochromat 20×/1.0 DIC objective.

### Western Blot

Protein lysates were prepared from cultured cells by lysis in RIPA buffer (ThermoFisher) supplemented with protease and phosphatase inhibitor cocktails (Roche) followed by sonication. Protein concentrations were determined using a BCA assay (ThermoFisher). Equal amounts of protein were denatured in Laemmli sample buffer, separated by Nu-PAGE 4-12% Bis-Tris gels (Invitrogen), and transferred to Nitrocellulose membranes (Bio-Rad). Membranes were blocked with SuperBlock (ThermoFisher) for 1 h at room temperature, then incubated overnight at 4 °C with primary antibodies against IRF3 (D38B9, Cell Signaling, 4302T), p-IRF3 (Ser396 4D4G, Cell Signaling, 4947T), RIG-1 (D14G6, Cell Signaling, 3743T), p-eIF2S1 (Abcam, ab32157), eIF2S1 (Atlas Antibody, HPA064885), and alpha-tubulin (DM1A, Millipore, 05-829) at manufacturer-recommended dilutions. After washing, membranes were incubated with HRP-conjugated secondary antibodies (Abcam) for 1 h at room temperature. Signals were detected by chemiluminescence (SuperSignal West Femto, ThermoFisher) and imaged using a ChemiDoc system (Bio-Rad).

### Agarose Gel Assay

RNA products generated by in vitro transcription were analyzed by agarose gel electrophoresis. Briefly, 500 ng of RNA per sample was mixed with 6x loading dye and resolved on a 1% agarose gel prepared in 1x TAE buffer containing SYBR Safe (Invitrogen). Gels were run at 100 V for 30-40 min, and RNA integrity was assessed by band size under UV illumination using a ChemiDoc system (Bio-Rad).

### Transfection Efficiency and Intensity Quantification

Quantification of nuclear mScarlet3 fluorescence intensity and cell segmentation was performed using CellProfiler. Nuclei were identified using the Minimum Cross-Entropy thresholding method, optimized for the signal-to-background characteristics of the images. The expected nuclear size range was defined based on the known pixel dimensions to ensure accurate segmentation across all images. Shape-based declumping algorithms were applied to separate adjacent or overlapping nuclei, allowing precise single-cell measurements. The mean fluorescence intensity per nucleus was then extracted and used to calculate transfection efficiency and relative expression levels across experimental conditions.

### Zebrafish Maintenance

Zebrafish were maintained at 28.5 °C at the UC Berkeley Fish Facility in accordance with an approved IACUC protocol (AUP-2019-09-12560-2; last approved December 16, 2025) and at the Janelia Research Campus under a separate IACUC protocol (#25-0278). Embryos were obtained from Casper zebrafish (mitfa^w2/w2^; mpv17^a9/a9^), which were selected for their optical transparency^36^.

### Embryo Microinjections

Injection solutions contained 240, 480, 600, or 720 pg of mRNA, 1 µL of injection buffer (0.1 M KCl, 0.1 percent phenol red salt, 0.1 mM EDTA, 1 mM Tris, pH 7), and RNase-free water to a final volume of 5 µL. One-cell embryos were injected directly into the cell with 3 nL per embryo at an injection speed of 10 nL/s using a Nanoject III Programmable Nanoliter Injector (Drummond Scientific).

For four-color zebrafish injections, solutions containing 150 pg each of mRNAs encoding mTagBFP2SKL, 4xCOX8-mStayGold, LAMP1-mScarlet3, and HaloTag-Sec61β were prepared in the same manner. One-cell embryos were injected into the cell with two sequential 0.5 nL injections using an Eppendorf FemtoJet 4i.

### Embryo Care

Embryos were kept in Danieau Buffer after injections and kept at 28.6 ° C, all dead or non-viable embryos were removed at 5 and 24 hours post injection. Embryos used in imaging were kept in a 6-well plate while all other embryos were kept in glass Pyrex dishes.

### Imaging Zebrafish

Zebrafish were immobilized using 3X Tricaine buffer (75 mg tricaine powder; 0.75 mL 0.3M Tris, pH 8.0; 0.075 mL NaOH; 73.35 mL 1X Danieau buffer^37^ . Imaging was performed on a Nikon SMZ18 with an ORCA-Flash4.0 C11440 (Hammamatsu) in MicroManger.

Four-color imaging was performed in 2 dpf zebrafish expressing mTagBFP2SKL, 4×COX8-mStayGold, LAMP1-mScarlet3, and HaloTag-Sec61β. HaloTag was labeled overnight with 4 µM JF635-HaloTag ligand. Embryos were dechorionated, anesthetized using 1X Tricain, mounted in capillary tubes, and imaged using a MuVi-SPIM.

### Image Analysis

Image analysis was performed using python3 and the BioImage package, the code for this analysis can be found at this link (https://github.com/coralnh/HC_mRNA_project). To account for camera and background noise, the average background intensity was acquired by capturing three empty fields of view in both acquisition channels (blue and green LEDs) and calculating the average intensity. This value was subtracted from each respective channel and then the average intensity per frame was calculated. As each embryo received the same dose of dual mRNAs (control MCP-mStaygold and test condition of MCP-mScarlet3) the background corrected red channel was normalized to the background corrected green channel to account for injection variability throughout the different conditions.

### Danionella cerebrum Maintenance, Microinjection, and imaging

*Danionella cerebrum* were maintained at Janelia Research Campus in accordance with an approved IACUC protocol (23-0244.27). Embryos were obtained from *mitfa* (-/-), which were selected for their optical transparency^38^. At the single-cell stage, embryos were microinjected with 1 nL of a solution containing 150 pg each of mRNAs encoding mTagBFP2-SKL, 4XCOX8-mStayGold, LAMP1-mScarlet3, and HaloTag-Sec61β. At 3 days post fertilization, larvae were labeled with JF635-HTL at a final concentration of 5 µM for 3 hours^39^, anesthetized with 1X Tricaine, and transferred to a CellVis 18-well coverglass chamber. Imaging was performed on a Zeiss LSM 980 equipped with Airyscan using CO-8Y Multiplex mode and a Plan-Apochromat 20X/0.8 NA M27 objective. Acquired images were processed as maximum intensity projections.

### Stronylocentrotus purpuratus (purple sea urchin) Maintenance, Microinjection, and imaging

Adult *Strongylocentrotus purpuratus* were collected and shipped from Point Loma Marine Invertebrate Lab (Lakeside, CA). Adult sea urchins are kept at 14°C in tanks with artificial seawater made from Instant Ocean^©^. Sea urchin males and females were spawned by shaking or via intracoelomic injection of 0.5 to 1 mL of 0.5 M KCl. Embryos were incubated at 14°C and cultured in filtered natural seawater collected from Indian River Inlet (University of Delaware, Newark, DE).

Microinjections were performed as previously described with modifications^40^. A vertical needle puller PL-10 (Narishige International) was used to pull the injection needles (1 mm glass capillaries with filaments) (World Precision Instruments). *mTagBFP2-SKL, 4x-COX8-mStayGold, LAMP1-mScarlet3* and *Halo-Sec61*β mRNA were lyophilized and resuspended with RNase-free water. The injection solution contains 2.25 µg of each mRNA, 20% sterile glycerol, in 3µL. Approximately 1-2 pL of solutions were injected into each newly fertilized egg. Injections were performed using the Pneumatic PicoPump with a vacuum (World Precision Instruments).

Embryos at morula (18 hpf), blastulae (24 hpf), gastrula (48 hpf), and larval stages (72 hpf) were collected and fixed with 4% paraformaldehyde (Electron Microscopy Sciences) in filtered sea water for 15 min at room temperature and washed 3 times 10 min with PBST (Tween-20 0.01%). The embryos were incubated with 500 nM JF646HTL^39^ for 1 hour. Imaging was performed on a Zeiss LSM 980 equipped with Airyscan with an EC Plan-Neofluar 40x/1.30 Oil DIC M27 objective.

