## Supplementary Information for "*mRNAbow*: A versatile gene expression system for multiplexed fluorescent imaging using optimized *in vitro* transcribed mRNA"


I. **Detailed protocol for IVT mRNA Production**
   a. **Background Information**
      i. **5' Untranslated Regions**
      ii. **Open Reading Frame (Translon)**
      iii. **3' Untranslated Regions**
   b. **DNA Template Preparation**
   c. ***In vitro* transcription**
   d. **RNA Purification**

II. **Supplementary Table 1.** List of Plasmids

III. **Supplementary Table 2.** qPCR primers

###  I. Detailed protocol for IVT mRNA Production

#### a. Background Information

##### i. 5’ Untranslated Regions (5’ UTRs)

The 5’ untranslated region (5’ UTR) is a non-coding sequence upstream of the open reading frame (ORF) that encodes the protein of interest. It plays a central role in regulating translation initiation, stability, and localization of mRNA[1]. A key element within the 5’ UTR is the 5’ cap structure, a hallmark of eukaryotic mRNAs that mediates recruitment of the translation machinery. The cap consists of a 7-methylguanosine (m7G) linked via a 5’-5’ triphosphate bridge to the first transcribed nucleotide (**Fig. 1**). This structure is recognized by the eukaryotic initiation factor 4E (eIF4E), which, together with eIF4G and eIF4A, forms the eIF4F complex. The eIF4F complex facilitates ribosome recruitment by guiding the 40S ribosomal subunit to the 5’ end of the mRNA and promoting scanning for the start codon[2].

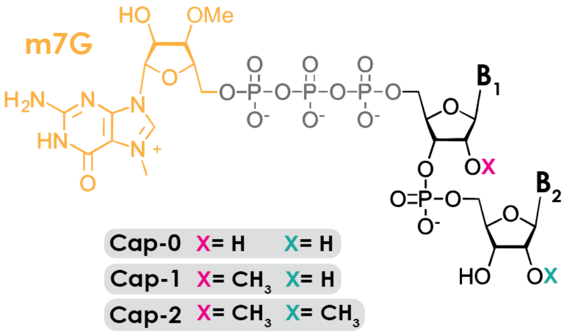


**Fig. 1.** Structure of 5’ cap structure, including Cap-0, Cap-1, and Cap-2.

Cap modifications modulate both immune recognition and translational efficiency[3]. In the Cap-0 structure, only the guanosine cap is methylated. Cap-1 adds a 2’-O-methyl group to the first ribose sugar, while Cap-2 includes 2’-O-methylations on both the first and second riboses. In mammalian systems, Cap-1 markedly reduces innate immune activation by preventing RIG-I recognition[4], whereas Cap-2 arises through gradual cytosolic methylation of Cap-1 and varies among cell types[5]. The Cap-1 structure enhanced protein expression compared to Cap-0[3]. CleanCap M6, which yields a Cap-1 structure containing an N6-methyladenosine at the first transcribed base, further improves protein production by stabilizing the cap and inhibiting decapping of exogenously delivered mRNAs[6]. In this study, we used CleanCap M6.

In addition to the cap structure, the 5’ UTR itself contributes to proper ribosome positioning at the correct start codon[7]. Because mammalian ribosomes scan from the 5’ end of the mRNA and tend to initiate translation at the first AUG encountered[2], the presence of upstream AUGs within the 5’ UTR can lead to translation from alternative reading frames and thereby reduce expression of the intended protein. The efficiency of initiation also depends on the Kozak sequence[8], a consensus motif located immediately upstream of the start codon that optimizes recognition by the ribosomal machinery. This sequence varies among species, so its modification may be necessary to enhance expression in systems other than mammalian cells.

In vaccine development, the 5’ untranslated region (5’ UTR) has been extensively optimized to improve mRNA stability, translation efficiency, and fidelity[9]. In this study, the 5’ UTR sequence employed corresponds to that used in the BNT162b2 (Pfizer-BioNTech COVID-19) vaccine. The sequence was reported by Andrew Fire and colleagues and made publicly available via his GitHub repository (https://github.com/NAalytics/Assemblies-of-putative-SARS-CoV2-spike-encoding-mRNA-sequences-for-vaccines-BNT-162b2-and-mRNA-1273), as shown below where red indicates the putative Kozak sequence:

5'- GAGAATAAACTAGTATTCTTCTGGTCCCCACAGACTCAGAGAGAAC**CCGCCACC**- 3'

##### ii. Open Reading Frame ("Translon")

The open reading frame (ORF) or translon encodes the protein of interest and represents the core functional element of the mRNA construct. Optimization of the ORF sequence is crucial in efficient protein production, as codon usage, GC content, and secondary structure can strongly influence translation efficiency and protein yield[10]. Codon optimization strategies replace rare codons with synonymous codons preferred in the target expression system, thereby improving ribosomal processivity and reducing premature termination. Additionally, adjusting GC content and minimizing stable secondary structures near the start codon can enhance ribosome scanning and initiation. There are several commercially available online software to perform codon optimization, including GenScript, VectorBuilder, or IDT.

The ORF should begin with a canonical start codon (ATG) that defines the correct reading frame. This start codon is typically embedded within an optimized Kozak consensus sequence to ensure precise initiation by the 40S ribosomal subunit. The coding region should terminate with one of the three standard stop codons (UAA, UAG, or UGA). Both Moderna mRNA-1273 and BioNTech/Pfizer BNT-162b2 used multiple stop codons which reduce the likelihood of translational readthrough and unwanted C-terminal extensions[9].

##### iii. 3' Untranslated Regions (3' UTRs)

Similar to 5' UTR, 3' untranslated region (3' UTR) is a non-coding sequence downstream of the open reading frame (ORF) that plays a central role in regulating mRNA stability, localization, and translational efficiency[11]. The 3' UTR interacts with a variety of RNA-binding proteins and microRNAs that modulate mRNA degradation rates and influence subcellular distribution[11]. In eukaryotic systems, the presence of a well-defined 3' UTR and a poly(A) tail enhances translation efficiency by facilitating the formation of the closed-loop complex, in which poly(A)-binding protein (PABP) interacts with eIF4G at the 5' cap to promote ribosome recycling[2].

In mRNA therapeutics and vaccines, the 3' UTR has been extensively optimized to maximize expression and stability[9]. Empirical screening has identified specific 3' UTRs like those derived from human α-globin, β-globin, or viral sequences that prolong cytoplasmic mRNA half-life and support high protein output[9].

The poly(A) tail downstream of the 3' UTR is equally important for stability and translational activation. In mammalian IVT mRNAs, optimal poly(A) tail lengths typically range from 100 to 150 nucleotides[12]. However, long homopolymeric tracts are prone to instability when directly encoded in plasmid DNA, often leading to deletions or recombination during bacterial propagation[13]. To overcome this, most IVT systems adopt one of several strategies: using a moderately sized encoded poly(A) stretch (50-70 nt), embedding the poly(A) sequence through extension primers, or combining a shorter encoded tail with *in vitro* enzymatic extension using *E. coli* poly(A) polymerase to achieve the desired length. This hybrid design maintains plasmid stability while producing a fully functional, efficiently translated mRNA transcript.

In this study, the 3' UTR sequence was adapted from the mRNA-1273 (Moderna COVID-19) vaccine construct, as reported by Andrew Fire and colleagues (see above). This region was selected for its proven ability to enhance translational efficiency and stabilize mRNA transcripts in mammalian cells. To balance plasmid stability and transcript functionality, the 3' UTR is followed by an encoded stretch of approximately 50 adenosine residues. This length proved sufficient in our experiments; however, if extended mRNA stability or half-life is desired, the poly(A) tail can be lengthened enzymatically or through extension PCR to achieve longer tail variants. Below is the our 3' UTR sequence:

**5'-**
GCTGGAGCCTCGGTGGCCTAGCTTCTTGCCCCTTGGGCCTCCCCCCAGCCCCTC
CTCCCCTTCCTGCACCCGTACCCCCGTGGTCTTTGAATAAAGTCTGAGTGGGCGG
CA - **3'**

#### b. DNA Template Preparation

This study employs T7 RNA polymerase, an enzyme derived from the *T7* bacteriophage that exhibits strong promoter specificity, transcribing only sequences located downstream of a T7 promoter. As outlined above, the essential elements for *in vitro* transcription using T7 polymerase include the T7 promoter, 5' untranslated region (5' UTR), gene of interest (GOI), 3' untranslated region (3' UTR), and a poly(A) tail. The plasmids designed for this system have been deposited in Addgene (see **Supplementary Table 1**) and are compatible with gene replacement using Gibson Assembly or In-Fusion Cloning, allowing flexible substitution of the GOI. The construct incorporates the 5' UTR from BNT162b2 and the 3' UTR from mRNA-1273, a combination previously reported to yield high stability and translational efficiency[14]. A compatible In-Fusion ready cloning kit following the same design is also commercially available from Takara. Additionally, one may order complete linear DNA fragments encoding all of these elements, providing a cloning-free alternative for direct use in *in vitro* transcription workflows.

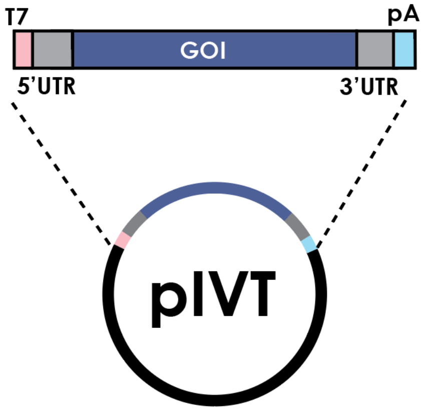


**Fig. 2** Template Plasmid Design. The essential element here is T7 promoter (pink), 5' UTR (grey), Gene of Interest (GOI), 3'UTR (grey), and polyA

For *in vitro* transcription, a linear DNA template is typically required, as circular plasmids can lead to rolling-circle transcription, producing overextended RNA transcripts lacking proper termination. While linearization is commonly achieved through restriction enzyme digestion, this approach can introduce complications such as incomplete digestion, star activity (nonspecific cleavage), and contamination with RNases or residual salts that inhibit transcription efficiency. Furthermore, depending on the enzyme and site used, blunt or overhang ends may cause heterogeneity at the RNA 3' terminus, which can affect translation fidelity and stability.

To overcome these limitations, previous studies have shown that adding two 2'-O-methyl (2'-O-Me) RNA nucleotides to the ends of PCR primers effectively terminates T7-driven transcription, preventing readthrough and the formation of double-stranded RNA by-products[15]. Based on this principle, we generated linear DNA templates by PCR amplification, using one primer encoding the T7 promoter at the 5' end and the other encoding a poly(T) tail with two 2'-O-Me RNA bases at its 5' terminus. This design produces clean, precisely terminated linear templates suitable for high-yield and high-fidelity transcription without the need for enzymatic digestion. PCR amplification was performed using Q5 Hot Start DNA polymerase with an annealing temperature of 64 °C, which produced a robust, high-fidelity linear DNA product optimized for subsequent *in vitro* transcription. It is important to check that the amplified DNA matches the expected size to ensure correct template generation and is free of other bands. If expected band is formed, perform PCR clean up using Monarch PCR/DNA Cleanup Kit.

**Forward Primer IVT** - 5'- CGAAAT**TAATACGACTCACTATAAG**GAATAA - 3'
**Reverse Primer IVT** - 5'- **mGmC**(T)$_{44}$ -3'

#### c. *In vitro* Transcription

For *in vitro* transcription (IVT), we employed Takara's PrimeCap T7 RNA Polymerase, a modified version of the *T7* bacteriophage RNA polymerase that enables co-transcriptional capping while minimizing the formation of double-stranded RNA (dsRNA) by-products. Conventional T7 RNA polymerase can produce antisense and self-complementary transcripts that anneal into dsRNA species, which are potent activators of innate immune sensors such as RIG-I, MDA5, and PKR[16,17]. These contaminants can reduce translational efficiency and increase cytotoxicity in transfected cells. In our hand, the PrimeCap T7 polymerase incorporates optimized buffer conditions and cap analog compatibility that suppress these unwanted products, yielding a highly pure, single-stranded mRNA population suitable for mammalian expression.

The co-transcriptional capping reaction uses a CleanCap M6 from TriLink that produces an authentic Cap-1 structure at the 5' end of the transcript. This design mimics the natural eukaryotic mRNA cap, ensuring efficient recruitment of the eIF4F complex and enhancing translational initiation. Co-transcriptional capping eliminates the need for separate enzymatic capping steps, reduces 5' heterogeneity, and improves mRNA integrity and translational performance[6]. For reference, the conventional T7 promoter directs transcription to initiate with a guanine (G), whereas the CleanCap AG or M6 system requires that transcription begin with **AG**, meaning the first two transcribed nucleotides downstream of the promoter are **AG** instead of a single **G** (**Fig. 3**).

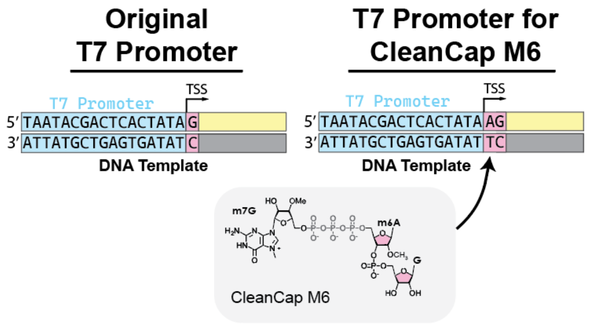


**Fig. 3** T7 promoter and start site design for CleanCap M6.

Typically, to further enhance transcript stability and reduce innate immune activation, modified nucleotides were incorporated into the IVT reaction. If wanted, N1-methylpseudouridine-5-Triphosphate (m1ΨTP) or 5-methoxyuridine-5-Triphosphate (5mo-UTP) can be used in place of UTP and 5-Methyl-Cytidine-5'-Triphposphate (5m-CTP) in place of CTP. These modifications reduce recognition by Toll-like receptors and cytosolic RNA sensors while enhancing translational efficiency in mammalian systems[18]. The combination of co-transcriptional capping and nucleoside modification results in transcripts with high stability, low immunogenicity, and superior protein expression in mammalian cells. Nonetheless, when dsRNA by-products are minimized, even unmodified nucleotides can support efficient protein production[15].

To minimize RNA degradation, we supplemented all reactions with SUPERase·In™ RNase Inhibitor (ThermoFisher). Work surfaces, pipettes, and centrifuge interiors were thoroughly decontaminated using RNase AWAY™ Surface Decontamination Solution, particularly when the centrifuge had previously been used for plasmid miniprep procedures, which often involve RNase A. All consumables, including pipette tips and tubes, were RNase-free, and non-DEPC-treated nuclease-free water was used in all reactions to prevent interference with enzyme activity while maintaining RNase-free conditions. Our procedure for running *in vitro* transcription reaction is the following:

| **Reagent** | **Volume** |
|---:|:---:|
| **SUPERase·In™ RNase Inhibitor (20 U/µL)** | 1 µL |
| **Nuclease-Free Water** | 5.4-x µL |
| **10X Transcription Buffer** | 2 µL |
| **100 mM ATP** | 2 µL |
| **100 mM CTP (or 5mCTP)** | 2 µL |
| **100 mM GTP** | 2 µL |
| **100 mM UTP (m1ΨTP or 5mo-UTP)** | 2 µL |
| **CleanCap Reagent M6** | 1.6 µL |
| **Template DNA (300 ng - 1 µg)** | x µL |
| **PrimeCap T7 RNA Polymerase** | 2 µL |
| **Total Volume:** | 20 µL |

The transcription reaction was incubated at 37 °C for 2 hours, during which high-yield RNA synthesis typically rendered the solution turbid or opaque. Subsequently, 8 units of RNase-free DNase I (NEB) were added directly to the reaction mixture, and the reaction was incubated for an additional 15 minutes at 37 °C to enzymatically degrade residual DNA templates. This 20 µL reaction generally yields 50 - 100 µg of mRNAs. Some modifications such as construct **7** (m1Ψ + 5mC) showed lower yield at our hands.

#### d. RNA purification

Following transcription, the RNA was purified to remove enzymes, unincorporated nucleotides, and other reaction components. Purification was performed using either the NEB Monarch RNA Clean-up Kit or by LiCl precipitation, both of which yielded comparable results. The Monarch kit provides rapid column-based purification, while LiCl precipitation selectively recovers long RNA transcripts and removes shorter abortive species. These steps effectively eliminate residual proteins, DNA, and small molecules, producing high-purity RNA suitable for downstream transfection or formulation. The purified mRNA was typically eluted in 80 µL of RNase-free water, and its concentration was determined by NanoDrop spectrophotometry. RNA integrity was assessed by agarose gel electrophoresis to confirm transcript size and quality. The purified mRNA was diluted to a final concentration of 200 ng/µL in RNAse free water, aliquoted into 10-20 µL volumes in RNase-free PCR tubes, and stored at -80 °C for long-term preservation. Under these conditions, the mRNA remained stable for at least three months. Once thawed, samples were not refrozen, as repeated freeze-thaw cycles can promote hydrolysis and fragmentation of RNA. Instead, thawed aliquots were stored at 4 °C and used within two weeks to maintain transcript integrity and translational efficiency.

#### II. **Supplementary Table 1.** List of IVT plasmids generated for this study

| Nucleus |
|---|
| MCP-Halo |
| MCP-mScarlet3 |
| MCP-mStayGold |
| MCP-mTagBFP2 |
| --- |
| mTagBFP2-HP1a |
| mCerulean3-HP1a |
| mStayGold-HP1a |
| mScarlet3-HP1a |
| Halo-HP1a |
| --- |
| H2B-mTagBFP2 |
| H2B-mCerulean3 |
| H2B-mStayGold |
| H2B-mScarlet3 |

| Lysosome |
|---|
| LAMP1-Halo |
| LAMP1-mScarlet3 |
| LAMP1-mStayGold |
| LAMP1-moxCerulean3 |

| Endoplasmic Reticulum |
|---|
| Halo-Sec61b |
| mScarlet3-Sec61b |
| mStayGold-Sec61b |
| mEmerald-Sec61b |
| moxCerulean3-Sec61b |

| Actin |
|---|
| Lifeact-Halo |
| Lifeact-mScarlet3 |
| Lifeact-mStayGold |
| Lifeact-mTagBFP2 |
| Lifeact-moxCerulean3 |

| Intermediate Filament |
|---|
| Vimentin-Halo |
| Vimentin-mScarlet3 |
| Vimentin-mStayGold |
| Vimentin-mTagBFP2 |
| Vimentin-moxCerulean3 |

| Mitochondria |
|---|
| 4xCOX8-mStayGold |
| --- |
| TOMM20-mTagBFP2 |
| TOMM20-moxCerulean3 |
| TOMM20-mStayGold |
| TOMM20-mScarlet3 |

| Microtubule |
|---|
| Ensconsin-Halo |
| Ensconsin-mScarlet3 |
| Ensconsin-mStayGold |
| Ensconsin-mTagBFP2 |
| Ensconsin-moxCerulean3 |
| --- |
| EB3-mScarlet3 |
| EB3-mStayGold |
| EB3-mTagBFP2 |
| EB3-moxCerulean3 |

| Golgi |
|---|
| SiT-mTagBFP2 |
| SiT-moxCerulean3 |
| SiT-mScarlet3 |
| SiT-mStayGold |
| SiT-Halo |
| --- |
| GalT-mTagBFP2 |
| GalT-moxCerulean3 |
| GalT-mStayGold |
| GalT-mScarlet3 |
| GalT-Halo |

| Empty Vector / Control |
|---|
| POI-Halo-IVT |
| POI-mScarlet3-IVT |
| POI-mStayGold-IVT |
| POI-mTagBFP2-IVT |
| POI-moxCerulean3-IVT |

| Calcium Sensor |
|---|
| WhaloCaMP1a |
| WhaloCaMP1b |
| jRCaMP1a |
| jRCaMP1b |
| GCaMP8s |
| GCaMP8m |
| GCaMP8f |
| GCaMP7f |

| Peroxisome |
|---|
| mTagBFP2-SKL |

| Endosomes |
|---|
| mStayGold-Rab5a |
| mTagBFP2-Rab5a |
| mScarlet3-Rab7a |
| mStayGold-Rab7a |
| mTagBFP2-Rab7a |
| mScarlet3-Rab11a |
| mStayGold-Rab11a |
| mTagBFP2-Rab11a |

| Phase Separation |
|---|
| mTagBFP2-DCP1 |
| moxCerulean3-DCP1 |
| mScarlet3-DCP1 |
| Halo-DCP1 |
| --- |
| G3BP1-mTagBFP2 |
| G3BP1-moxCerulean3 |
| G3BP1-mScarlet3 |
| G3BP1-Halo |

| Plasma Membrane |
|---|
| Myr-mTagBFP2 |
| Myr-moxCerulean3 |
| Myr-mStayGold |
| Myr-mScarlet3 |
| Myr-Halo |

#### III. Supplementary Table 2. qPCR primers

| qPCR Primer sets | |
|---|---|
| ***β-Actin (ActB)*** | |
| Forward Set 1 | CAC CAT TGG CAA TGA GCG GTT C |
| Reverse Set 1 | AGG TCT TTG CGG ATG TCC ACG T |
| ***Nuclear mScarlet3*** | |
| Forward Set 1 | GCT GAA TGG ATC AGC TCT AAC |
| Reverse Set 1 | CCT TTA GGC ACC TCG ACT TT |
| Forward Set 2 | CGA CTT CAA GAC CAC CTA CAA |
| Reverse Set 2 | CTG GTG ATG TCC AGC TTT CT |
