## Supplementary figures and images for "*mRNAbow*: A versatile gene expression system for multiplexed fluorescent imaging using optimized *in vitro* transcribed mRNA"

### Supplementary Figure 1

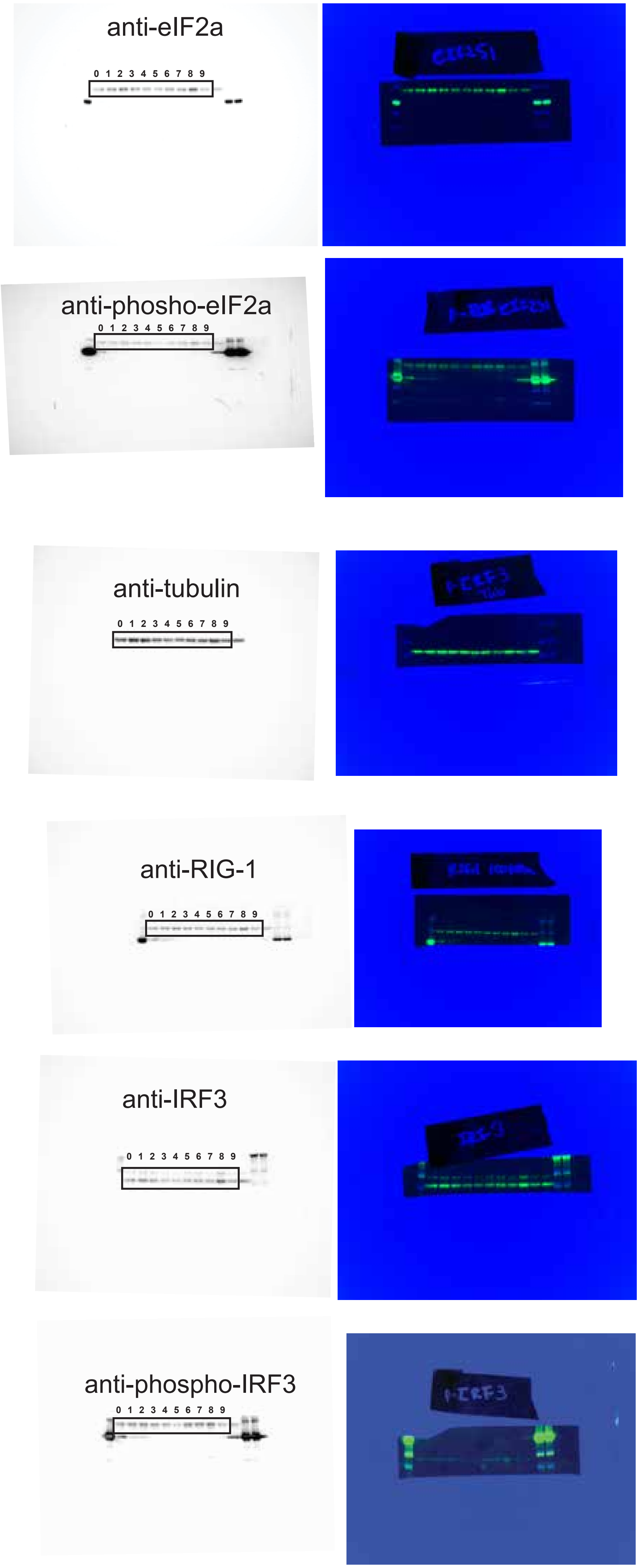


Supplementary Fig. 1

### Supplementary Figure 2

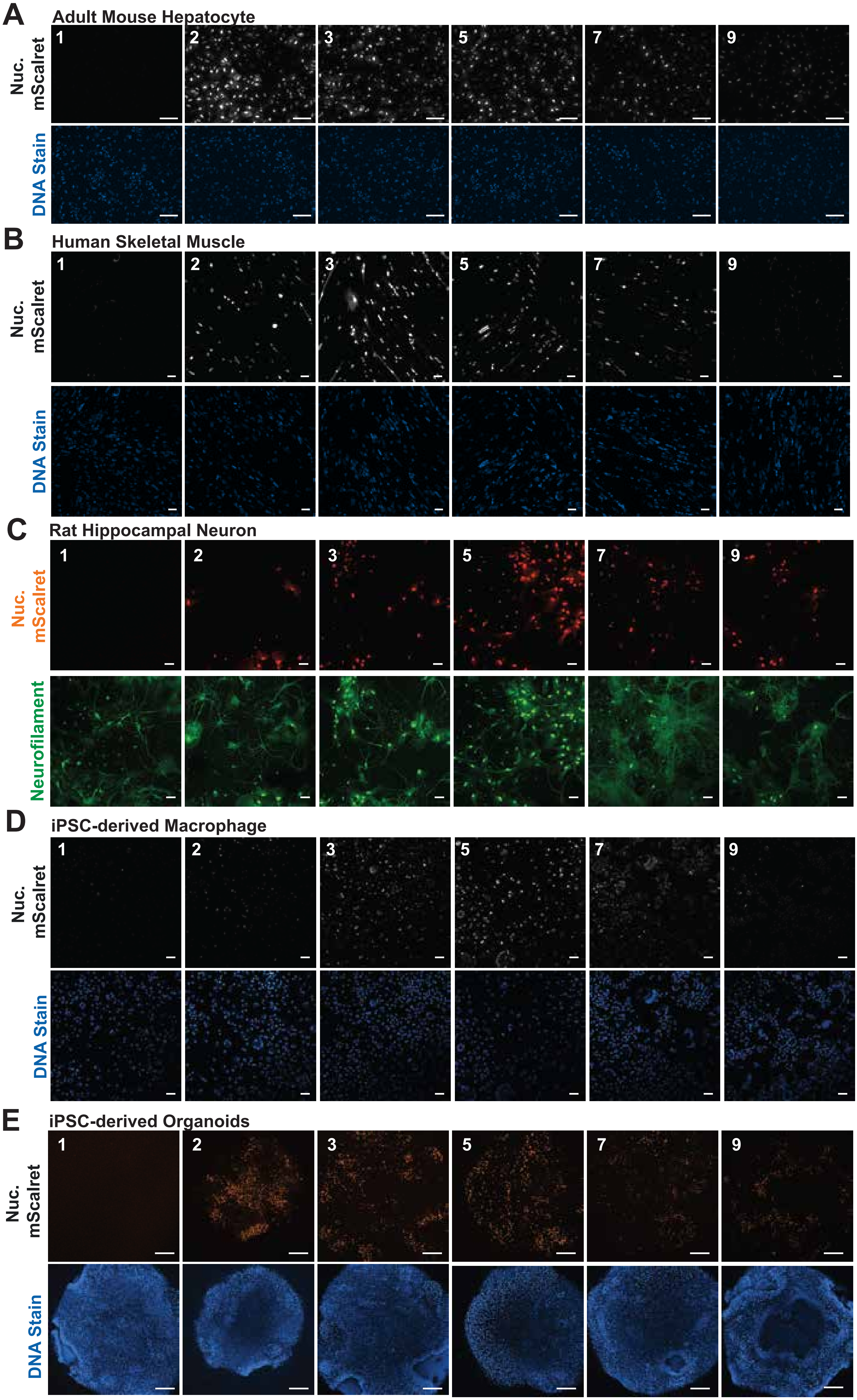


Supplementary Fig. 2

### Supplementary Figure 3

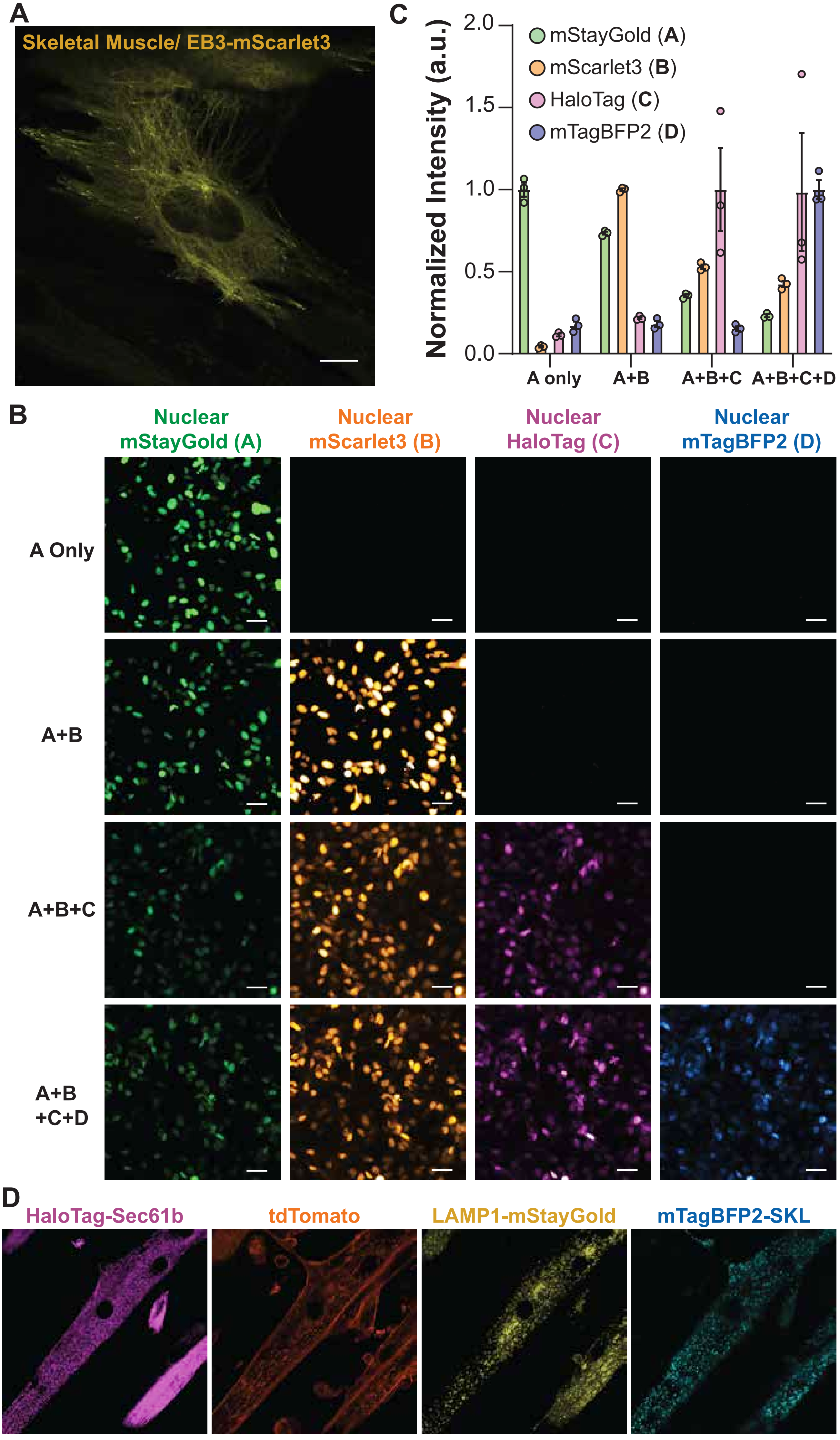


Supplementary Fig. 3

### Supplementary Figure 4

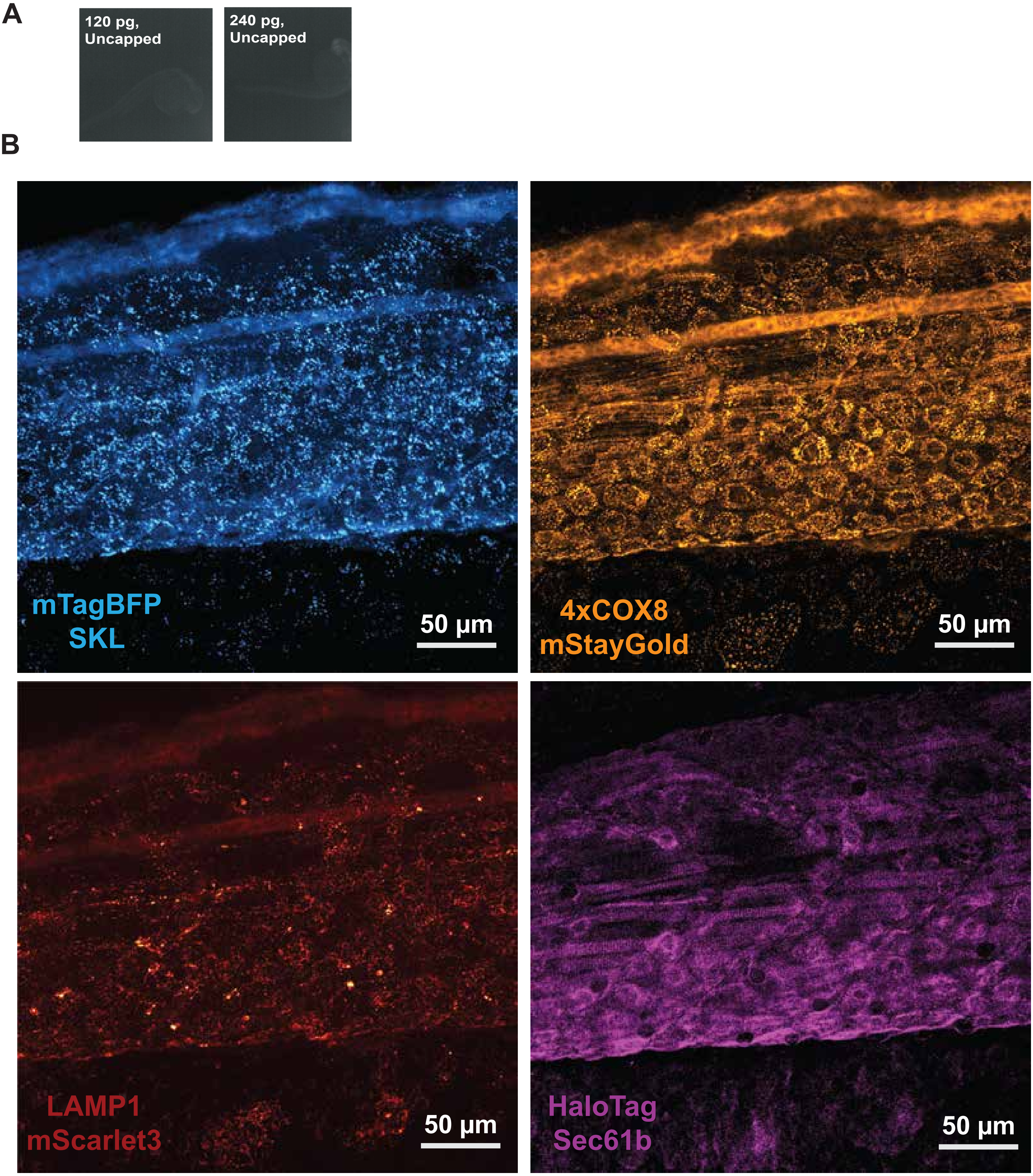


Supplementary Fig. 4

### Supplementary Figure 5

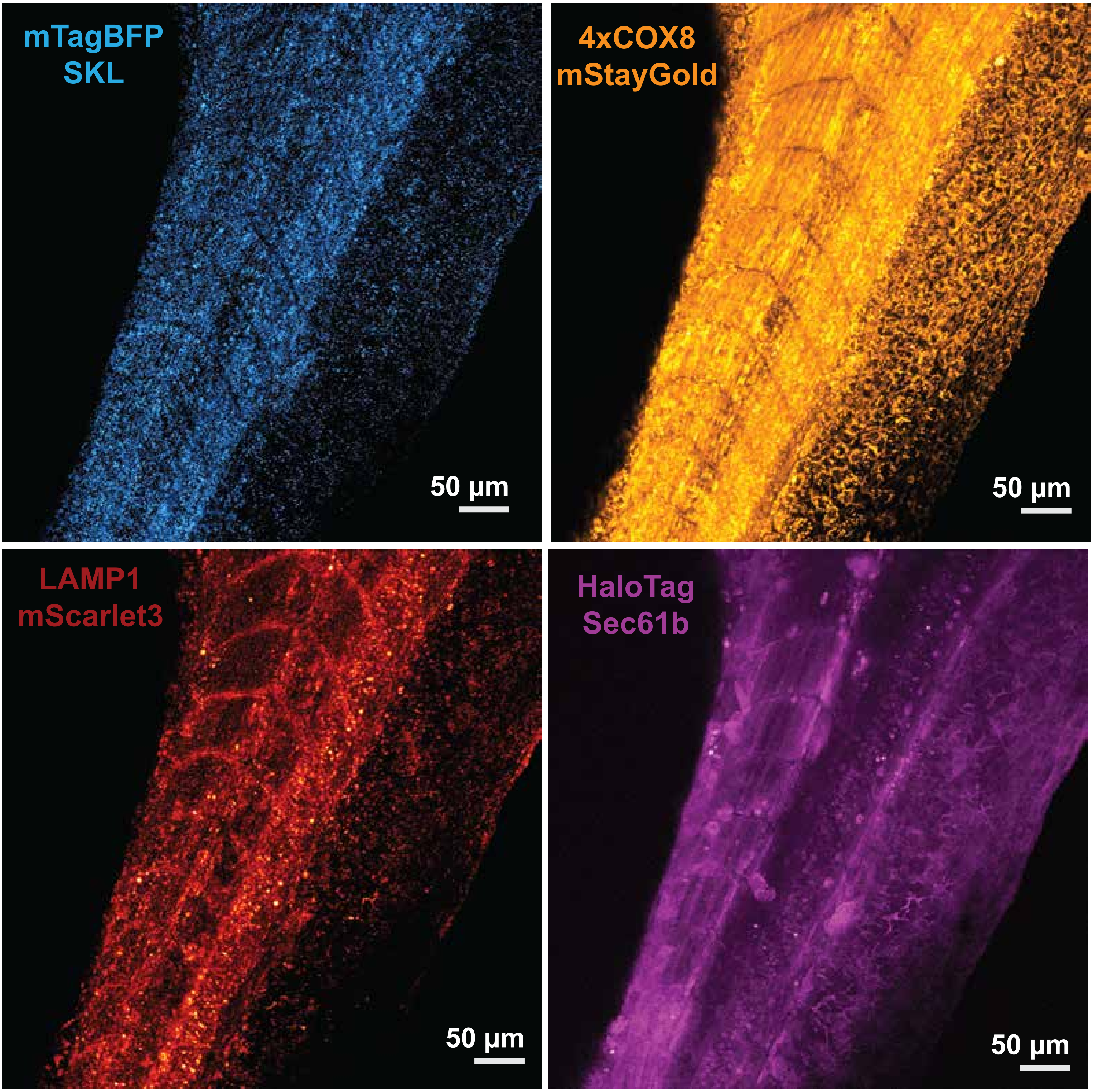


**Supplementary Fig. 5**

### Supplementary Figure 6

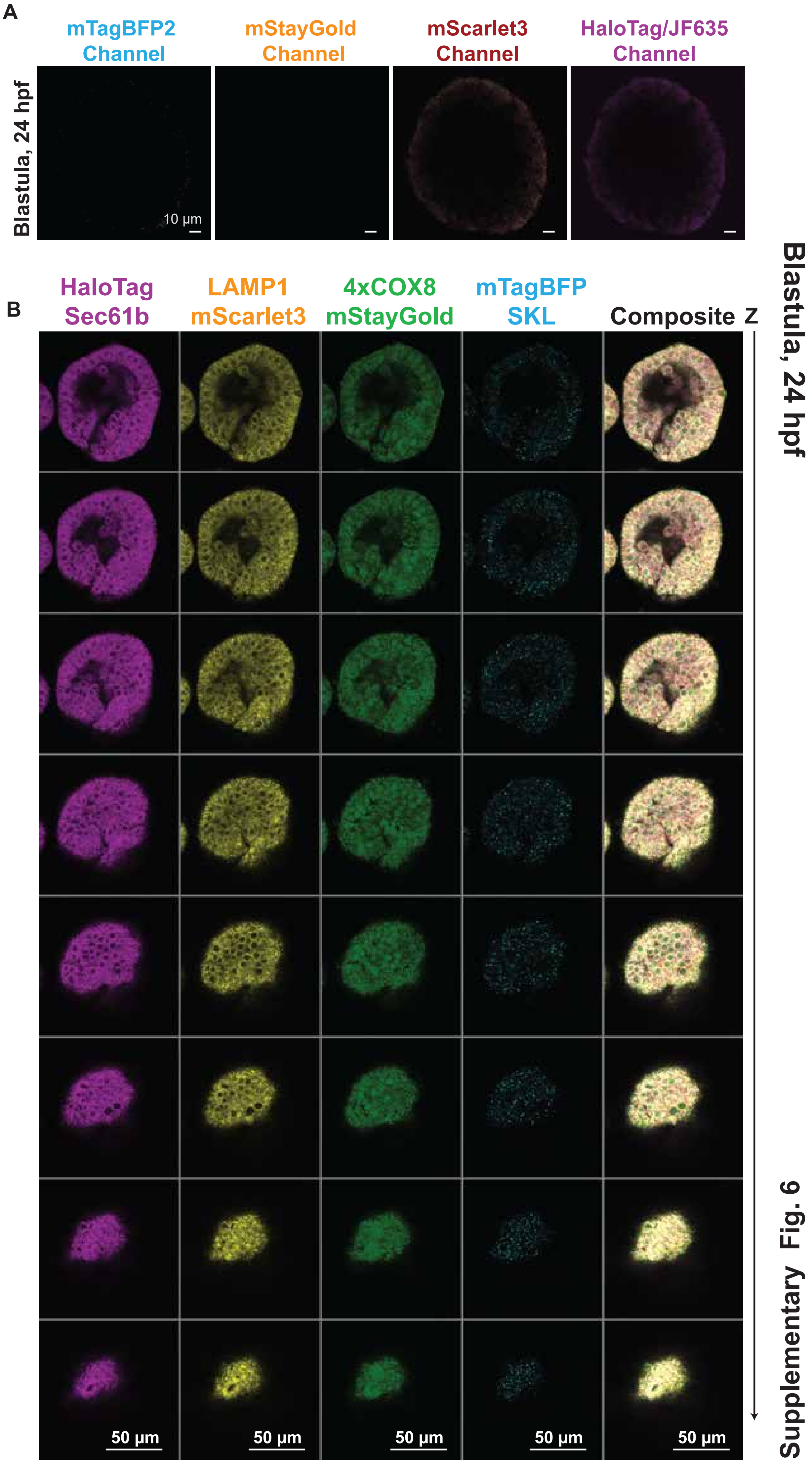


Supplementary Fig. 6

### Supplementary Figure 7

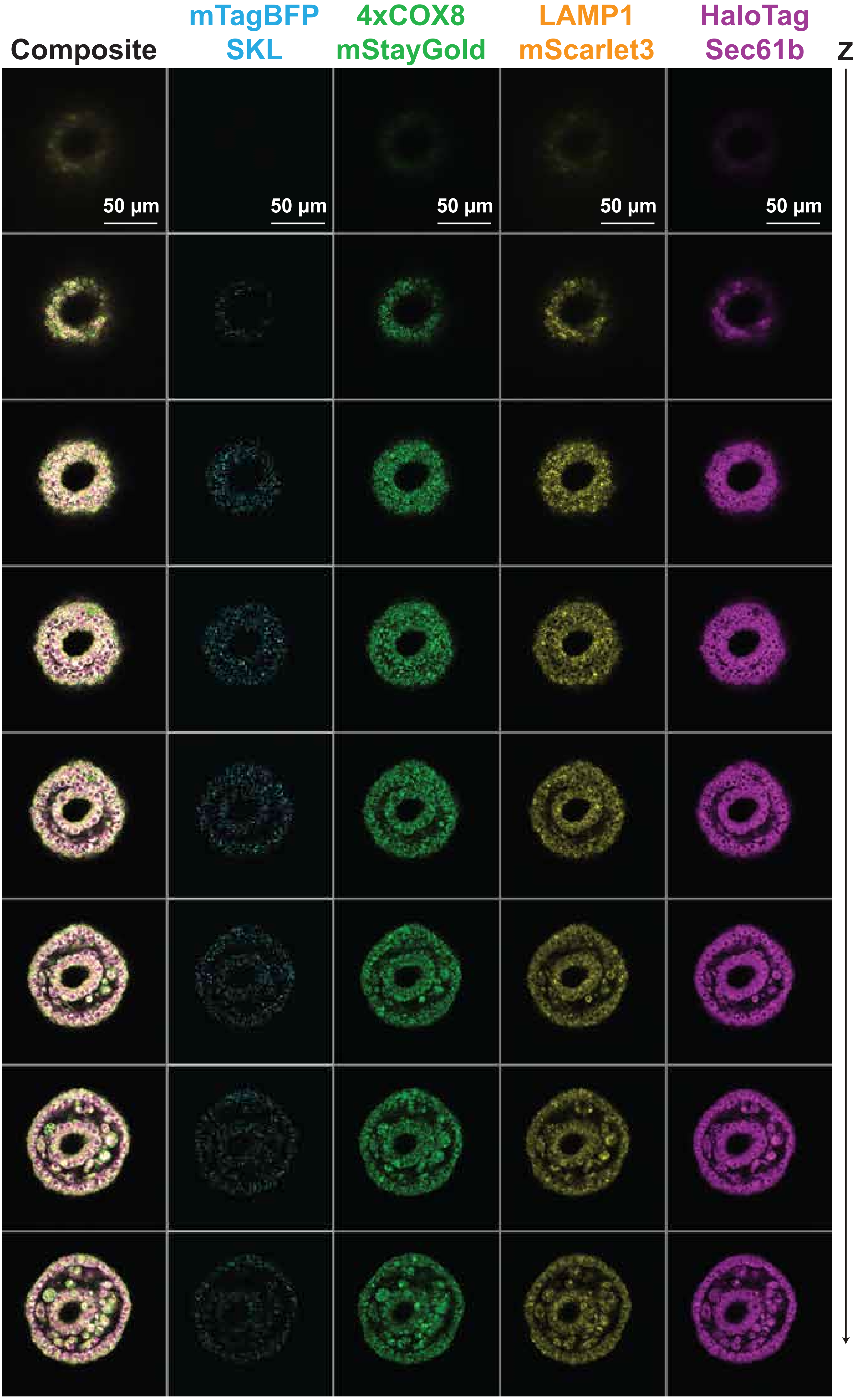


Supplementary Fig. 7

### Supplementary Figure 8

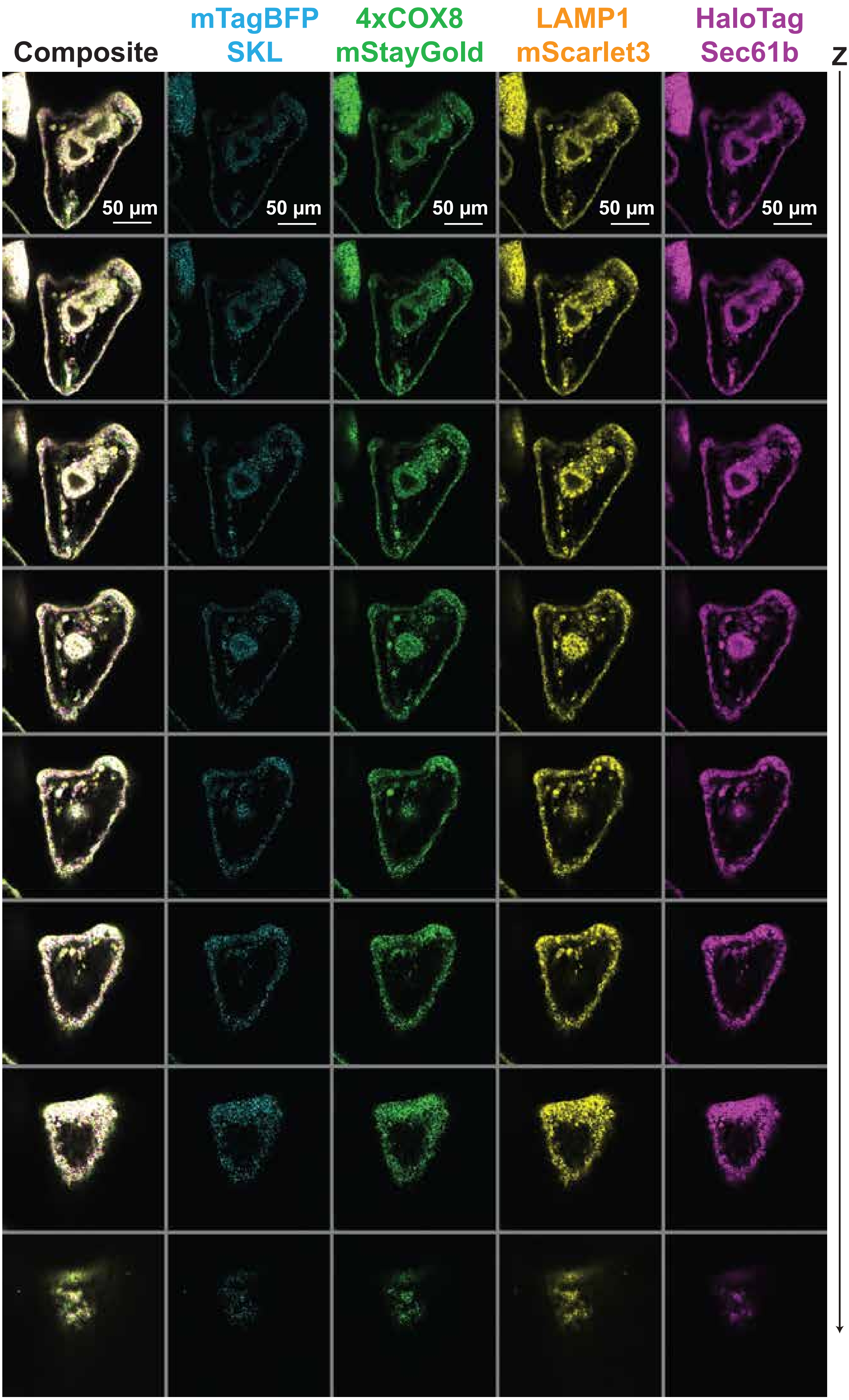


Supplementary Fig. 8
